# Circulating neutrophils from patients with early breast cancer have distinct subtype-dependent phenotypes

**DOI:** 10.1101/2023.04.19.537022

**Authors:** Anisha Ramessur, Bana Ambasager, Iker Valle Aramburu, Freddie Peakman, Kelly Gleason, Christoph Lehmann, Venizelos Papayannopoulos, Raoul Charles Coombes, Ilaria Malanchi

## Abstract

**Purpose:** A high number of circulating neutrophils is a poor prognostic factor for breast cancer, where evidence of bone marrow cancer-dependent priming is found. However, how early this priming is detectable remains unclear.

**Patients and Methods:** Here, we investigate changes in circulating neutrophils from newly diagnosed breast cancer patients before any therapeutic interventions. To do this, we assessed their lifespan and their broader intracellular kinase network activation states by using the Pamgene Kinome assay which measures the activity of neutrophil kinases.

**Results:** We found sub-type specific L-selectin (CD62L) changes in circulating neutrophils as well as perturbations in their overall global kinase activity. Strikingly, breast cancer patients of different subtypes (HR+, HER2+, triple negative) exhibited distinct neutrophil kinase activity patterns indicating that quantifiable perturbations can be detected in circulating neutrophils from early breast cancer patients, that are sensitive to both hormonal and HER-2 status. We also detected an increase in neutrophils lifespan in cancer patients, independently of tumour subtype.

**Conclusions:** Our results suggest that the tumour-specific kinase activation patterns in circulating neutrophils may be used in conjunction with other markers to identify patients with cancer from those harbouring only benign lesions of the breast. Given the important role neutrophil in breast cancer progression, the significance of this sub-type of specific priming warrants further investigation.

**Clinical Relevance:** The current study aims to investigate cancer-specific changes in circulating neutrophils in patients with newly diagnosed early breast cancer before any therapeutic intervention. We found L-selectin (CD62L) changes in circulating neutrophils from patients with early-stage breast cancer compared to healthy volunteers, which is an indication of an early phenotypical change. Moreover, these changes in CD62L were dependent on the breast cancer sub-type, showing opposing trends according to the hormonal receptor status of the tumour. Importantly, this subtype dependent phenotypic alteration was reflected in broader intracellular signalling perturbation when measuring intracellular kinase activity. Moreover, those cancer perturbed neutrophils, show expanded life span when cultured ex vivo, suggesting an alteration in their physiologic state. The tumour-specific kinase activation patterns in circulating neutrophils may be useful in conjunction with other markers to distinguish patients with cancer from those with benign lesions of the breast.

## Introduction

Recently neutrophils have become a subject of intensive research in cancer due to the growing evidence of their involvement in various aspects of cancer onset and progression^1^. Tumour infiltrating neutrophils have a variety of different pro-tumoral activities, ranging from immunosuppression to directly influencing tumour growth and metastatic activity. Neutrophils have been the subject of recent reviews highlighting how their pro-tumour functions might be related to their physiologic tissue protection functions^2^ as well as their potential for therapeutic targets^2, 3^. In addition to their role in the tumour and metastatic environment, there is evidence for the ability of a tumour to mobilize and perturb neutrophils systemically^3^. Indeed, systemic changes in circulating neutrophils has been found in patients across a range of different cancer types^4^. These changes have been shown to have repercussions for neutrophils’ tumour-promoting functions in metastases^5–7^. Moreover, studies using high dimensional analysis such as cytometry by time-of-flight mass spectrometry (CyTOF) or single cell analysis revealed the presence of different neutrophils phenotypes and signatures pointing to the existence of various neutrophil subsets^8, 9^. While the relevance and the cause of this neutrophil diversity are still to be understood, these phenomena point to their previously underestimated plasticity in response to environmental signals. Therefore, it is paramount to use the evidence about the different cellular phenotypic states acquired in the context of cancer to deepen our understanding of neutrophil biology as well as to help develop more effective therapeutic strategies for patients.

Breast cancer is the leading cause of cancer-related deaths among women worldwide, with an estimated 2.4Lmillion new cases and 523L000 deaths reported in 2015^10^. There are distinct subtypes of breast cancer: Hormone Receptor positive (HR+), whereby breast tumour growth is responsive to oestrogen or progesterone, and Hormone Receptor negative (HR-), where breast cancer growth is independent of such overactive hormonal signalling. Breast cancer can also present with a higher expression of human epidermal growth factor receptor 2 (HER2) on tumour cells which is termed HER2 positive (HER2+) breast cancer. HER2+ can be a variable in HR+ breast cancer (tumour type is defined either as HR+HER+ or HR+HER-) or HR-breast cancer (either HR-HER+, or HER2-, HR-which is termed triple negative (TN) breast cancer). A high neutrophil to lymphocyte ratio (NLR) has been shown to have a prognostic value for breast cancer patients^11^, especially for stage I patients, where a high NLR could stratify patients who developed distant metastasis, but not local recurrences^12^, suggesting that systemic changes related to neutrophils may occur early in the disease process and could be meaningful for metastatic progression.

In animal models of human breast cancer, tumour mediated granulopoiesis has been reported^13, 14^, and the systemically mobilized neutrophils show different phenotypic properties depending on the genetic driver of the breast cancer model used^15^. For instance, circulating neutrophils in patients with cancer as well as mouse models of multiple cancer types, showed increased expression of a fatty acid transporters supporting lipid metabolism^7^.

This cancer-driven granulopoiesis and neutrophil priming has consequences for metastatic progression, since this leads to the accumulation of neutrophils within distant organs, supporting the growth of subsequently disseminating cancer cells^13–15^. This is in keeping with the clinical evidence that a high NLR has prognostic value for patients with breast cancer developing distant metastasis^12^.

The current study aims to investigate cancer-specific changes in circulating neutrophils in patients with newly diagnosed early breast cancer (EBC) before any therapeutic intervention. We found L-selectin (CD62L) changes in circulating neutrophils from patients with EBC compared to healthy volunteers (HVs), which is an indication of an early phenotypical change. Moreover, these changes in CD62L were dependent on the breast cancer sub-type, showing opposing trends according to the hormonal receptor status of the tumour. Importantly, this subtype dependent phenotypic alteration was reflected in broader intracellular signalling perturbation when measuring intracellular kinase activity. Moreover, those cancer perturbed neutrophils, show expanded life span when cultured ex vivo, suggesting an alteration in their physiologic state. Most strikingly, these tumour-specific kinase activation patterns in circulating neutrophils may be used in conjunction with other markers to identify patients with cancer from those harbouring only benign lesions of the breast.

## Results

### Early Breast Cancer Patients show tumour HR status specific L-selectin changes circulating neutrophils

Cancer mobilised neutrophils have been reported to undergo significant phenotypic changes in patients across a range of cancers^4^. However, most of the studies investigate circulating neutrophils in cancer patients with more advanced disease and following treatment interventions.

In a mouse model of breast cancer, mice harbouring spontaneous MMTV-PyMT breast tumours that induced neutrophilia^14^, we found that systemically mobilised neutrophils showed reduced surface expression of L-selectin, (CD62L^low^) in various tissues (Supplementary Figure 1 and 2). In this proof-of-concept study, we aimed to test if similar changes in circulating neutrophils could be detected in newly diagnosed patients with EBC. To do this, we compared neutrophils from blood samples of 44 treatment-naive patients with EBC (after biopsy, but prior to surgery or neoadjuvant chemotherapy) with those from 44 paired sex and age matched healthy volunteers (HVs) collected on the same day as each breast cancer sample. Moreover, to accommodate for any confounding factors such as the effect of adrenaline or cortisol released by patients experiencing stress in anticipation of treatment, we recruited an additional group of patients who were undergoing surgery for benign breast disease. Selected patients’ data and selection criteria are listed in Supplementary Table 1-3. We firstly analysed circulating neutrophils from EBC patients and paired HVs by FACS to assess whether a similar change in CD62L could be observed in patients. In line with the neutrophilia detected in the animal model of EBC^14^, neutrophils were found to make up a higher proportion of total live cells in the blood (Figure 1a). These data reinforce the notion that neutrophils are systemically responding to the presence of breast cancer. When we analysed the percentage of CD62L^low^ neutrophils in patients and compared to HVs, we found that, consistent with the murine HR negative MMTV-PyMT breast cancer model (Supplementary Figure 2), neutrophils from HR-negative patients had significantly elevated proportion of CD62L^low^ subpopulations compared to HVs (Figure 1b). Conversely, HR-positive patients had significantly lower levels of CD62L^low^ subpopulations compared to matched HVs (Figure 1c). These data indicate that perturbations in the surface expression of CD62L were detectable very early in cancer and are dependent on breast cancer’s hormonal receptor status (Figure 1d, e).

**Figure 1:**
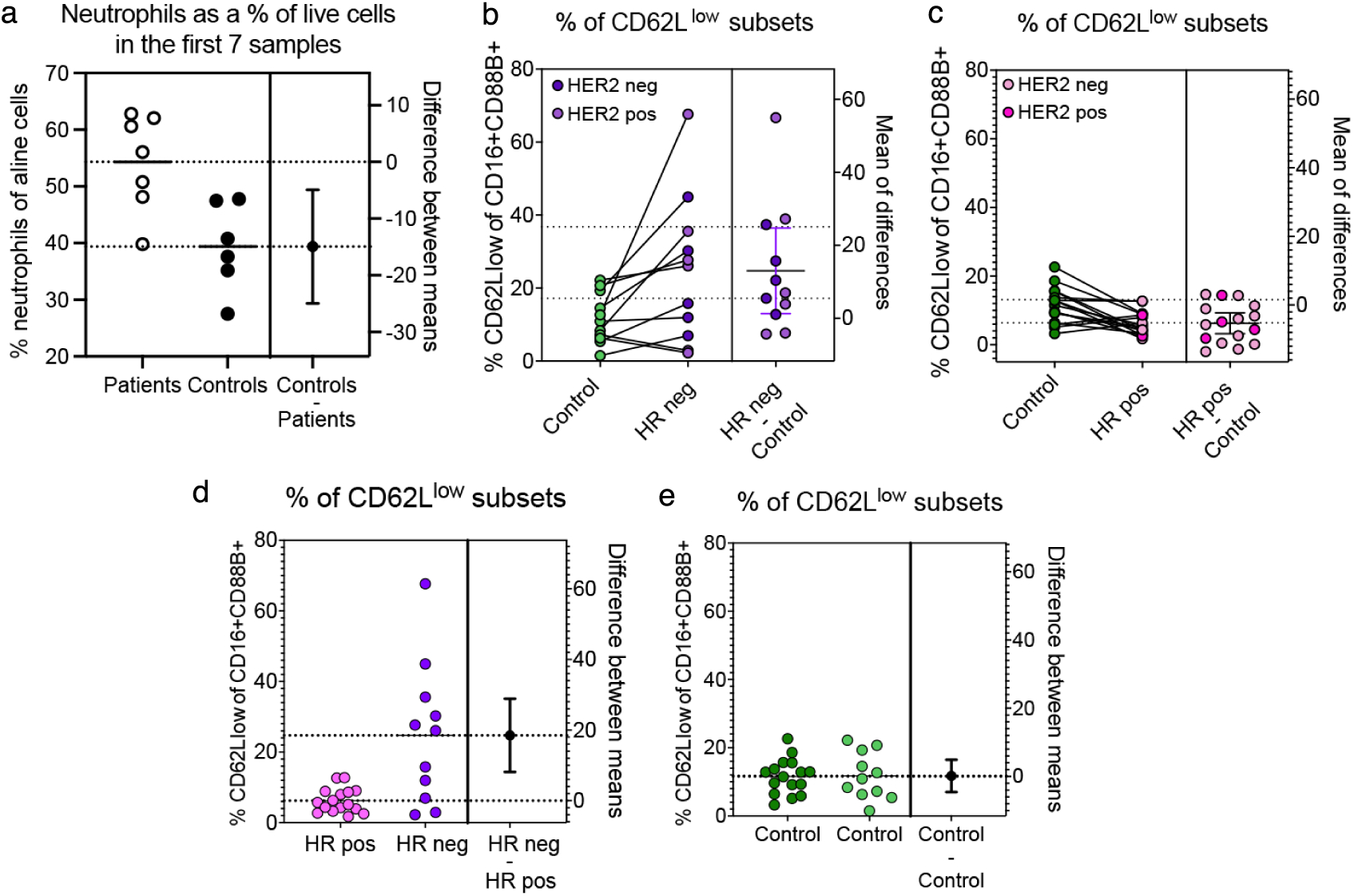
**a.** Flow cytometry analysis showing neutrophils (CD16+CD66B+) as a % of total live cells in the blood for 8 patients with early breast cancer compared to HVs. Statistical analysis by Unpaired T- test (two-tailed). Data are represented as mean ± standard error of the mean. **P = 0.0073. **b-e.** Flow cytometry analysis of CD62L^low^ neutrophil in patients with: **(b)** HR negative breast cancer compared to HVs, Statistical analysis by paired T-Test (two-tailed). P**= 0.0331; **(c)** HR positive breast cancer compared to HVs, Statistical analysis by paired T-Test (two-tailed). P**= 0.0019; **(d)** HR positive compared to HR negative breast cancer, Statistical analysis by unpaired T-Test (two-tailed). P**= 0.0011; **(e)** HVs of HR positive compared to HVs of HR negative breast cancer, Statistical analysis by unpaired T-Test (two-tailed). P value not significant (P= 0.9455). Data is represented as mean ± standard error of the mean.

### Circulating neutrophils from patients with early breast cancer show an increase lifespan

As we could detect differences in markers associated with neutrophils aging via flow cytometry in circulating neutrophils of patients with EBC, we next assessed if a difference in their properties could be detected in isolated neutrophils ex vivo. Neutrophils are reported to have a short lifespan; however, this can be influenced in steady state by different tissue context^17^ as well as by factors generated during inflammation^18, 19^. We therefore measured neutrophils’ lifespan, upon their isolation from the patients’ circulation. We calculated the half-life of neutrophils in culture for 32 hours and found that neutrophils from patients with EBC (Supplementary figure 3a-b), regardless of subtype, had a longer half-life compared to paired HVs when cultured in their own plasma (Figure 2a). Moreover, the fold increase in the half-life of neutrophils from EBC patients mildly correlated with the neutrophil to lymphocyte ratio (NLR) values (Supplementary figure 3c). In addition, when neutrophils from patients with EBC were cultured with plasma from the paired HVs we still observed a significant increase in half-life compared to neutrophils from HVs (Figure 2b). Similarly, a difference was not observed when cancer patient-derived neutrophils were cultured in their own plasma or in HV plasma (Figure 2c). This suggests that neutrophils are intrinsically primed to live longer when isolated from blood of cancer patients. Moreover, plasma from patients with EBC did not show any effect on neutrophils from HVs (Figure 2d, e), reinforcing the idea that a change in neutrophil behaviour originates from a more complex in vivo priming.

**Figure 2:**
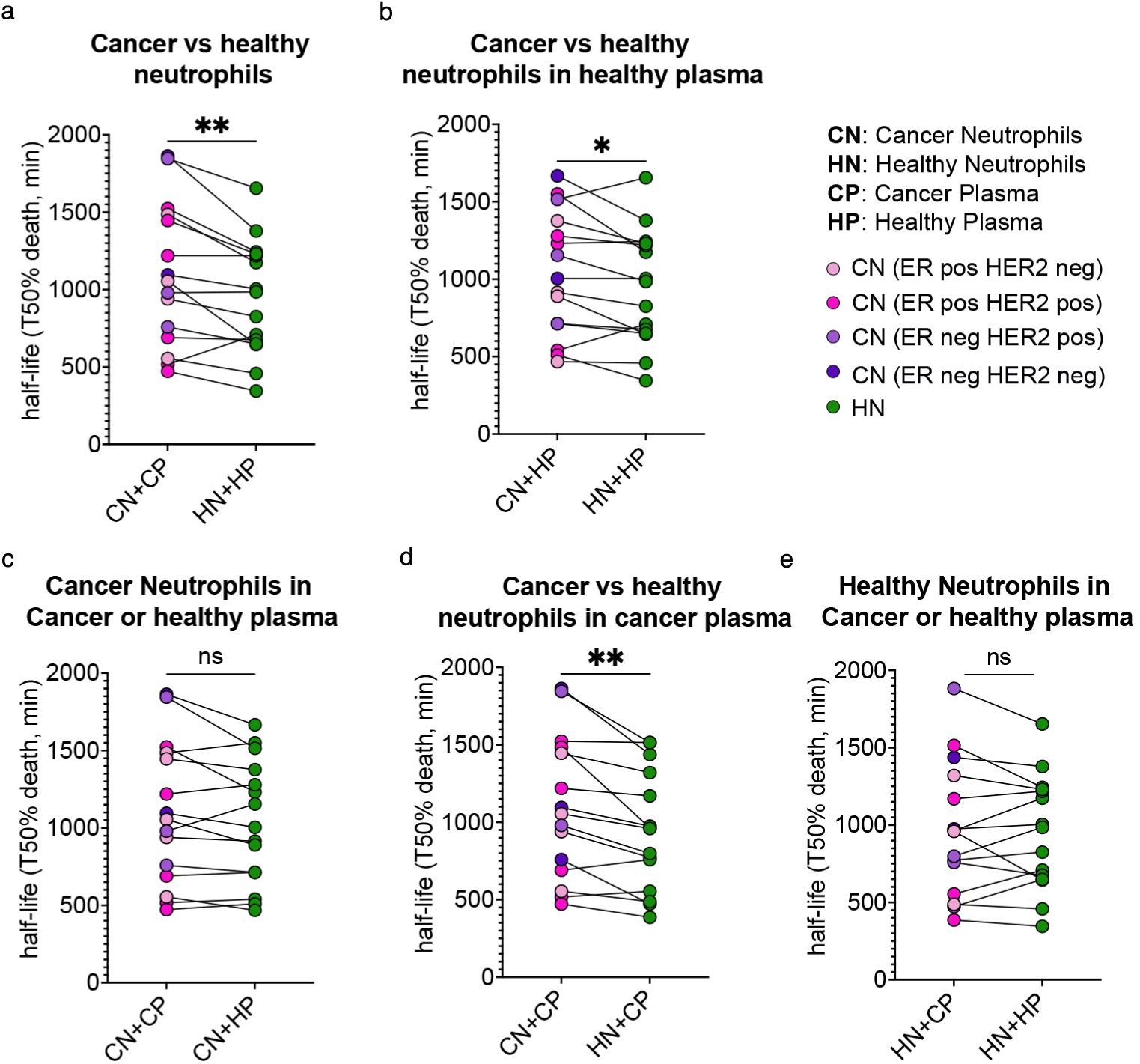
Functional assessment of neutrophil half-life as an indicator of lifespan for patients with breast cancer and HVs across a range of subtypes. **a.** Neutrophil half-life as measured by 50% death of neutrophils in minutes for neutrophils from cancer patients (CN) or paired HVs (HN) which are cultured in their own plasma (CP or HP respectively). Breast cancer subtypes are colour labelled. Statistical analysis using Wilcoxon matched pairs signed rank test. P* = 0.0052; **b.** Neutrophil half-life for neutrophils from cancer patients (CN) or HVs (HN) which are cultured in paired HV plasma (HP). Statistical analysis using Wilcoxon matched pairs signed rank test. P* = 0.0353; **c.** Neutrophil half-life for neutrophils from cancer patients (CN) which are cultured in their own plasma (CP) or paired HV plasma (HP). Statistical analysis using Wilcoxon matched pairs signed rank test. P value not significant (P=0.1205); **d.** Neutrophil half-life for neutrophils from cancer patients (CN) or HVs (HN) which are cultured in paired plasma from cancer patient (CP). Statistical analysis using Wilcoxon matched pairs signed rank test. P* = 0.0012; **e.** Neutrophil half-life for neutrophils from HVs (HN) which are cultured in their own plasma (HN) or plasma from paired cancer patients (CP). Statistical analysis using Wilcoxon matched pairs signed rank test. P value not significant (p=0.7197)

Taken together, we could detect an intrinsic priming affecting neutrophils in EBC patients which display a prolonged half-life.

### Circulating neutrophils in patient with early breast cancer showed perturbations in their overall intracellular kinase activity

Following the indications of an early perturbation detected by flow cytometry in circulating neutrophils from patients newly diagnosed breast cancer along with a prolongation of their lifespan, we aimed to assess if these phenotypic changes were reflected in appreciable overall signalling pathway activities. To do this we investigated their broader intracellular kinase network activation states using the Pamgene Kinome assay. To avoid unintentional activation due to antibody binding, circulating neutrophils were isolated from blood using a negative selection strategy. Total proteins were extracted from circulating neutrophils from EBC patients and their paired HVs and loaded onto kinome chips to functionally screen their kinases activity by quantitative measurements of phosphorylated target peptides specific to either serine/threonine or protein tyrosine kinases (STKs or PTKs respectively) (Supplementary Figure 4a and 4b). Specific peptides that increase or decrease output of phosphorylation can be linked to the kinase activity that typically recognised them. Collectively this analysis provides a functional readout of overall kinome activity in circulating neutrophils. Supplementary Figure 4c shows an example of the raw STK data represented as a heatmap for the HR-positive HER-2 negative group of patients. This heatmap represents the overview of phosphorylated peptides within neutrophils in patients and HVs samples prior to normalisation of the data. Only peptides which passed quality control were Log2 transformed, and combat corrected (normalised for batch and pair effect to account for experiments done on different days) prior to further analysis. The log fold change (LFC) of phosphorylated peptides between patients compared to paired HVs was calculated for each subtype of breast cancer (Supplementary Figure 4d, e). This showed there appears to be some variability between patients regarding the degree of phosphorylated peptides for individual patients compared to their paired HVs.

In order to identify global patterns in the data, we used Principal Component Analysis (PCA) to assess presence of clustering for the different subtypes of breast cancer compared to HVs for the two classes of kinases, STK and PTK family (Supplementary Figure 5). Overall, the results did not show obvious clustering of patient and HV groups, only in HR+ and triple negative patients some level of clustering could be observed compared to their paired HVs (Supplementary Figure 5b-c and 5f-g).

To get a better overview of the changes in kinase activity in the different breast cancer subtypes, we used Coral trees to plot all the kinases tested. Coral trees are used for visualization of the human kinome superfamily, representing both quantitative and qualitative data^16^. Using this analysis, the differences between the neutrophil kinome in all patients with a given breast cancer subtype compared to HVs became clearer (Figure 3). Interestingly, the entire neutrophil kinome, and particularly the PTK family, appears to be more perturbed in patients with early HR-positive compared to HR-negative breast cancer (Figure 3a). HR-negative disease is characterized by a milder increase in activity of the TK family compared to HVs while other kinase activity is overall reduced compared to the HVs (namely kinases within the CGMC family, AGC family and CK1 kinase) (Figure 3b). Strikingly, the presence of HER-2 positive disease appears to revert this dominant TK activation increase and induced a reduction compared to HVs in both HR-positive and HR-negative patients (Figure 3c, d). The rest of the kinome remains more activated compared to HVs.

**Figure 3:**
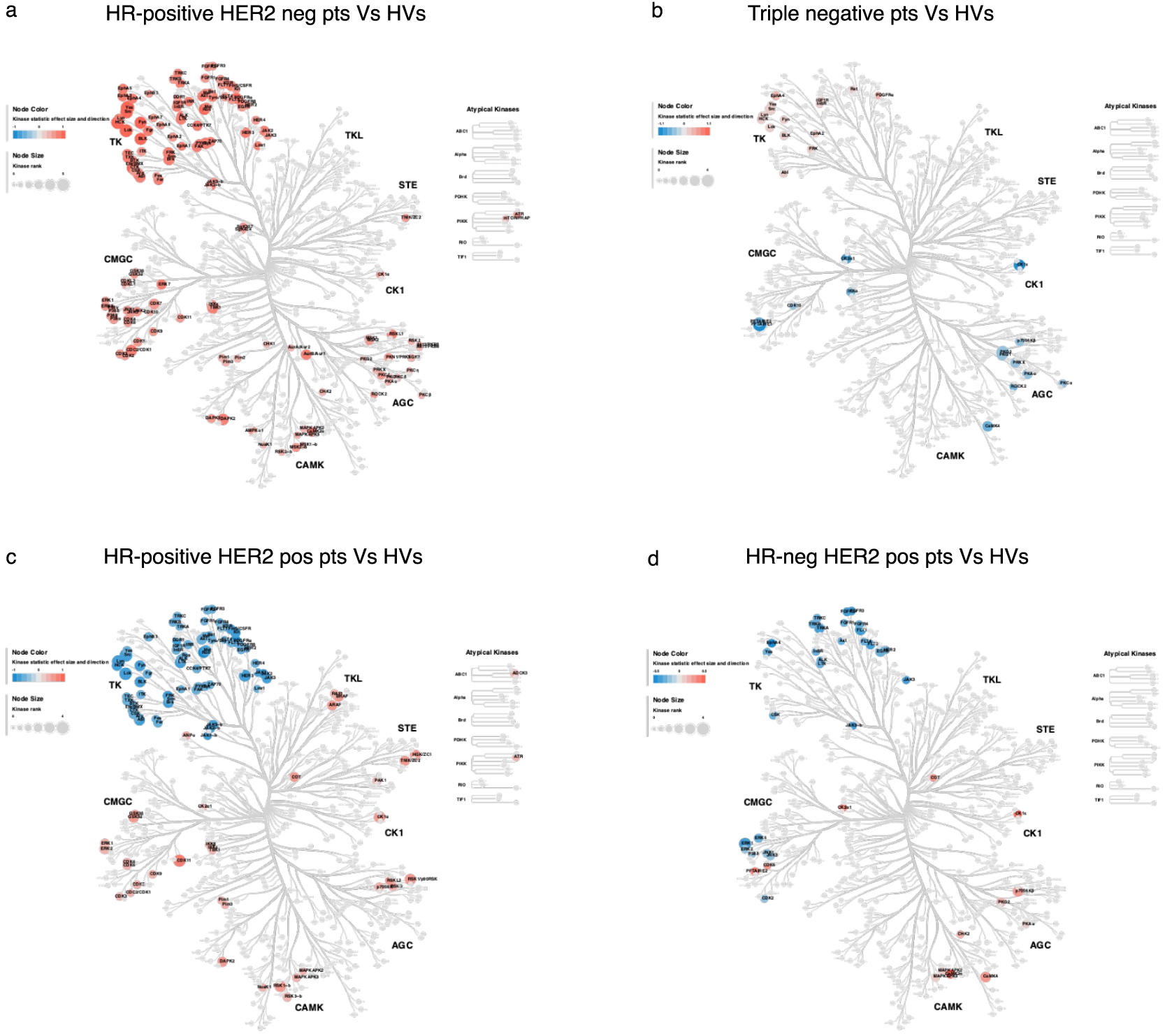
**a-d.** Coral trees to show the whole kinome family (including STK and PTK families) in the indicated breast cancer sub-types compared to HVs. Each coloured node represents a specific putative kinase which is upregulated (red) or down-regulated (blue) in activity in neutrophils in patients with breast cancer compared to HVs (each subtype of breast cancer compared to HVs is shown separately). Nodal size represents the median significance value for the individual kinase. **(a)** Coral tree to show differences in neutrophil kinome in patients with HR-positive HER2 negative breast cancer compared to HVs; **(b)** Coral tree to show differences in neutrophil kinome in patients with HR-positive HER2 positive breast cancer compared to HVs; **(c)** Coral tree to show differences in neutrophil kinome in patients with HR- negative HER2 positive breast cancer compared to HVs; **(d)** Coral tree to show differences in neutrophil kinome in patients with Triple negative breast cancer compared to HVs

Collectively these data show an early neutrophil cancer-specific priming in their overall intracellular kinome activity, that appears to be different accordingly to breast cancer subtype and deeply influenced by the presence of HER-2 positivity.

### The kinome activity of circulating neutrophils could possess the potential to discriminate cancer from patients with benign disease

As shown in Figure 3, the kinome activity signatures of circulating neutrophils from patients with breast cancer are perturbed compared to paired HVs and different accordingly to breast cancer subtype. However, there are differences between the EBC patients and HVs: only patients with cancer have previously undergone a biopsy and are under the stress of a surgery at the time of blood sampling. Therefore, to test if those differences are specific to cancer patients, we collected circulating neutrophils from patients that were diagnosed with benign breast disease after biopsy. Importantly, when analysing their kinome activity, while some perturbation could be observed, we did not detect alteration specifically in TK activity compared to paired HVs (Figure 4a). Given this data suggested a potential breast cancer specific change, we tested if the changes in kinome activity in circulating neutrophils could retain the potential to predict characteristics of the disease in patients, by performing two types of predictive model analysis. The raw data used for the analysis are provided in Supplementary Table 3 and included the two variables measured for each kinase activity by comparing each patient to its paired HV measured using the Pamgene platform. The two variables indicate the degree of significance for each kinase for that given patient vs paired HVs (MFS) and the direction of the change, either up or down regulated compared to HVs (KS) respectively (Supplementary File 1).

**Figure 4:**
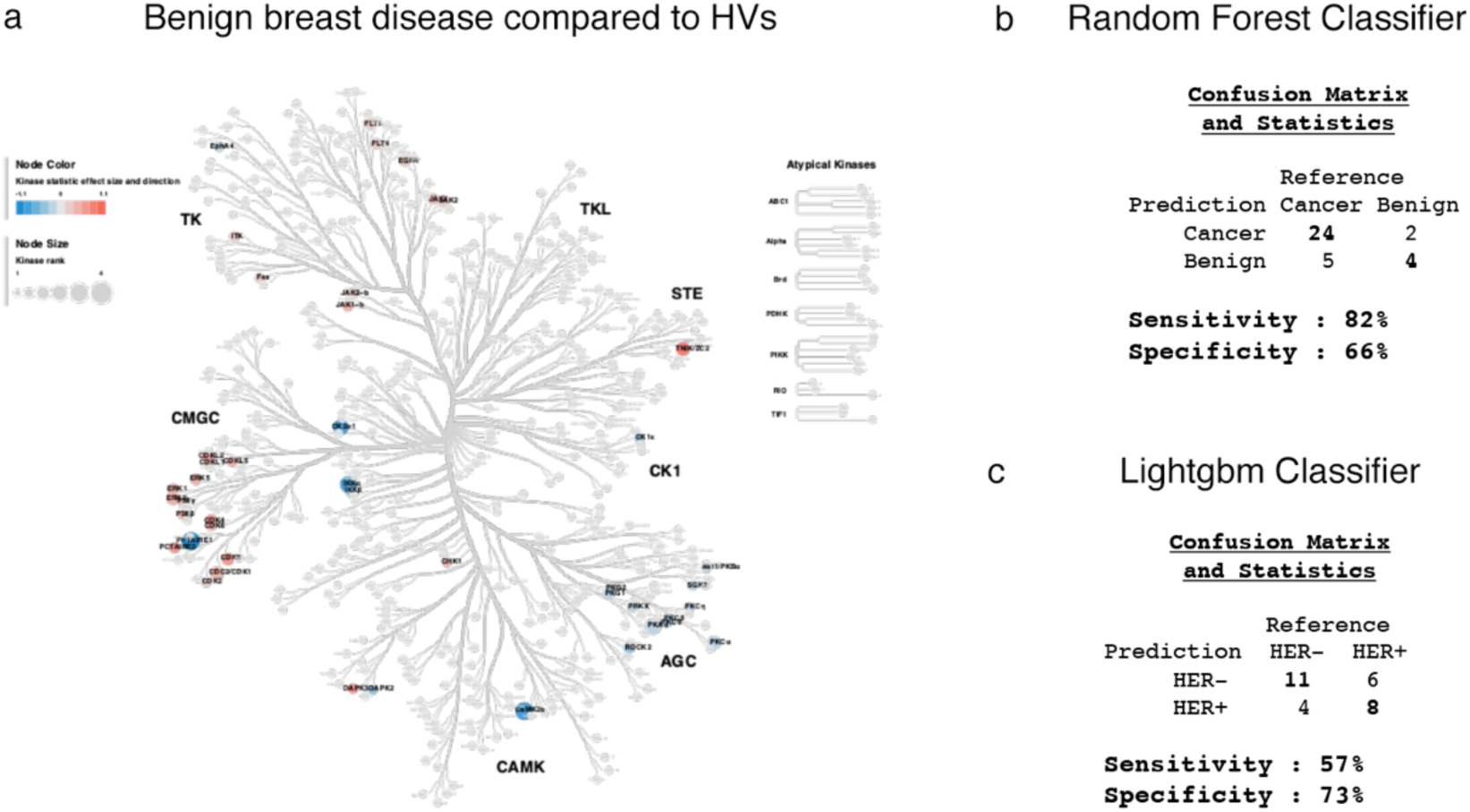
**a**. Coral trees to show the whole kinome family (including STK and PTK families) in neutrophil kinome in patients with benign breast disease compared to HVs. Each coloured node represents a specific putative kinase which is upregulated (red) or down-regulated (blue) in activity in neutrophils in patients with breast cancer compared to HVs (each subtype of breast cancer compared to HVs is shown separately). Nodal size represents the median significance value for the individual kinase. **b.** Outcome of predicted randomly selected patients for the benign or cancer group using a Random Forest classifier; **c.** Outcome of predicted randomly selected patients for the HER2 positive or HER2 negative cancer patients using a Lightgbm classifier.

Two machine learning algorithms were used for training the model according to the number of patients to compare in the two groups (Supplementary File 2). Firstly, we compared the 29 samples from patients of all types with the 6 samples from benign lesions. In this case we use a Random Forest algorithm to train the model, which is better suited for imbalanced data (a much fewer patient in one group). Strikingly, randomly selected patients which were not used to train the model, could be predicted to belong to either the cancer or the benign group with a sensitivity of 82% and a specificity of 66% (Figure 4b). Moreover, this model predicted that the kinase activity changes with the stronger weight were LCK and PKG1 (Supplementary Figure 6a). When looking at those two kinases in the Upstream Kinase Analysis (UKA) generated list of putative kinases based on peptides targeted for phosphorylation in patients with different breast cancer subtypes (Supplementary Figure 7), they are indeed among the kinases perturbed in all cancer patients but do not show changes in circulating neutrophils of benign lesions.

Secondly, given the marked effect of HER2 expression on PTK activity observed in patients of both breast cancer types compared to HV, we tested if HER-2 status could be predicted in cancer patients from the kinome perturbation in their circulating neutrophils. We compared the profiles of neutrophils isolated from both sub-types of breast cancer patients, either negative or positive for HER-2 mutation using Lightgbm algorithm to train the model since the number of patients in the two group are comparable. The results showed that we could identify the HER-2 status of patient with a specificity of 73% and a sensitivity of 57% (Figure 4c). Interesting the model predicted that the kinase activity change with the stronger weight was p70S6K beta (Supplementary Figure 6b).

In conclusion, by training a predictive model with only the few patients analysed here, we could show that the differences in tumour-induced kinome activity of circulating neutrophils could have the potential to distinguish between EBC and benign lesions and it could detect the presence of HER-2 mutation in breast cancer.

## Discussion

A large body of literature has drawn functional links between neutrophil activities and cancer onset and progression^20^. Most of these studies are performed in animal models, however recently, much evidence of the systemic perturbations of neutrophils in patients with cancer have been also described^4^. This led to the notion that neutrophils play a key role in cancer onset, progression, and resistance to therapy^1^. Moreover, certain genetic mutations can manifest as intrinsic properties of cancer which are known to shape the local immune landscape and the systemic engagement of neutrophils specifically^15, 21^.

Here we assessed firstly if systemic perturbations in circulating neutrophils could be detected very early in breast cancer and secondly, whether those perturbations are specific to the tumour and its sub-type. We selected patients diagnosed with different sub-types of EBC and analysed their circulating neutrophils before any treatment intervention. When isolated from the circulation, neutrophils can change and activate, thus introducing a high level of variability during handling. Therefore, a sex, age and menopausal status matched HV sample was also collected on the same day as the patient sample to be used in direct comparisons.

In a preclinical breast cancer model, previously shown to induce neutrophilia^14^, we detected an expansion of neutrophils with a reduction in L-selectin (CD62L^low^) a change that was previously reported in circulating neutrophils of Triple Negative (TN) breast cancer patients^22^. We found that in newly diagnosed cancer patients with TN breast cancer, the pool of CD62L^low^ neutrophils was also expanded, but our data showed that this change appears to be highly subtype specific. The results showed that neutrophils from HR+ breast cancer showed an opposite trend, where even fewer circulating CD62L^low^ neutrophils were present in patients with cancer compared to matched HVs. These data indicate that EBC already induces systemic changes in neutrophils, and that those changes might be influenced by the tumour’s receptor status.

Given this initial indication of a phenotypic change, we assessed if we could detect an intrinsic perturbation in circulating neutrophils of EBC patients. We performed a broad functional analysis of their kinase activities as an indicator of their intracellular signalling pathways. We used Pamgene Kinase analysis to assess differences between the activity of neutrophils’ lysates to phosphorylate kinase specific peptides when isolated from patients with EBC compared to paired HVs. The signal from the phosphorylation of each peptide was measured by the Pamgene machine and generated a prediction of the corresponding kinase responsible for the increase or decrease activity.

When assessing the PCA plots, the kinome data derived from the PTK family, could cluster neutrophils from patients and HVs. As kinome data derived from the STK family did not generate this separation, we concluded that patients with different subtypes of breast cancer compared to the paired HVs shows a more significant difference for the PTK family than the STK family.

When visualizing all kinase activities in Coral trees, it became more evident that the TK family of kinases are a hot spot of cancer-mediated perturbations. Interestingly, the HER-2 status of the patient’s tumour was a dominant factor in influencing the PTK activity. Even if with different extents, in both patients with HR-positive and HR- negative breast cancer, the presence of HER-2 expression in the tumour resulted in a strong downregulation in most of the TK family of kinases which would have otherwise been upregulated. The HR status of the patient’s tumour seemed to be more important than HER-2 status in influencing the STK activity in neutrophils.

The mechanism is uncertain but there is evidence that different myeloid infiltrates (neutrophils and macrophages) are present in the tumour microenvironment of different breast cancer type, which are important for cancer behaviour under treatment^23^, so there may be different soluble factors released by HR-positive and HR -negative tumour subtypes which differentially influence the intrinsic neutrophil kinome activity.

Surprisingly, the changes in neutrophil kinome for patients with the more aggressive subtype, Triple negative breast cancer compared to HVs are comparatively less than with other breast cancer subtypes. Nonetheless, it remains to be determined how the perturbation in overall kinase activity relates to neutrophils functions. We observed that neutrophil lifespan was prolonged independently of the type of kinome perturbations, suggesting that all EBC disease, regardless of the type of kinome perturbation, triggered a functional priming of neutrophils. When neutrophils from patients with breast cancer, independently of subtype, were cultured in their own plasma, we found an increase in lifespan compared to HV neutrophils (P= 0.0052). Plasma from patients with breast cancer did not extend the lifespan of HV neutrophils. Moreover, the increased lifespan of cancer-derived neutrophils remained significant when neutrophils from patients were cultured in plasma from HVs, but the effect was less striking. Collectively, these data suggest that the expansion in lifespan in circulating neutrophils from cancer patients was due to an intrinsic priming but was likely to be additionally supported by cancer-derived factors.

The increase in neutrophil half-life may in part explain the flow cytometry findings which showed patients with breast cancer had a higher proportion of circulatory neutrophils compared to HVs since neutrophils may survive longer and therefore accumulate systemically. Our data showing these early neutrophil alterations, are in line with the clinical observation of an increased neutrophil to lymphocyte ratio especially in the early disease^12^.

Most importantly, when analysing the neutrophil kinome of patients with benign breast disease, we detected low level of perturbations compared to patients with cancer. A prediction model trained on the kinome data of the patients analysed in this small pilot study, showed this to be sensitive and specific in predicting the presence of cancer. Taking into consideration the low number of patients investigated in this proof-of-concept study, our results suggest that a diagnostic blood test, based on the early cancer dependent perturbation in circulating neutrophils, could be developed. This would involve a larger kinome dataset obtained by the analysis of circulating neutrophils from larger patients’ cohorts with different EBC subtypes as well as benign disease patients, to generate models to reliably predict breast cancer at the same time as mammogram screening, even before histologic analysis derived by a formal biopsy.

Additionally, our results indicate that both hormonal status and the presence of HER-2 mutation profoundly influence the cancer dependent neutrophil priming, a phenomenon that warrants further mechanistic studies.

## Materials and Methods

### Statistical analysis

Unless otherwise stated, statistical analyses were done using Prism Version 9 (GraphPad software). Error bars for column plots represent average ± Standard error of the Mean (SEM).

When analysing the difference in one variable between two experimental groups, paired or unpaired student T-tests were carried out and p-values were obtained from this. One-way ANOVA was used for comparing the difference in one variable between two or more experimental groups. Two-way ANOVA was used to perform multiple comparisons or analyses between experimental groups. The significance level was set at p< 0.05 and the actual p values or symbols (*p < 0.05, ** p< 0.01, *** p< 0.001, **** p<0.0001) were used. Biological replicates are represented as n values.

#### Mouse strains and neutrophils quantification

The MMTV-PyMT mice in FVB were kept at the Biological Research Facility of the Francis Crick animal facilities and all animal procedures were performed in accordance with UK Home Office regulations under the Home Office project license PP5920580. The National Cancer Research Institute (NCRI) Guidelines for the Welfare and Use of Animals in Cancer Research were followed. When assessing primary tumour growth, a mean diameter of 1.5cm for single tumours was not exceeded. However, for multifocal disease such as MMTV-PyMT cancer, if there were no additional adverse welfare consequences for the animal, the total superficial tumour burden was allowed to exceed these dimensions when essential for the achievement of the scientific objective, namely spontaneous metastasis. Mice were monitored daily for signs of adverse effects.

MMTV-PyMT+ mice spontaneously developed a primary tumour and increased neutrophil presence in the blood, bone marrow, spleen and liver. Neutrophil infiltration was quantified by flow cytometry by anti-Ly6G antibody (clone 1A8 from BioXcell).

### Patient characteristics

Patients with EBC were selected based on HR status and HER2 status of their tumour. The term early breast cancer refers to the breast tumour being localised to the breast and includes patients with or without local lymph node involvement where the management is aimed for curative intent. The patient characteristics including age, menopausal status, medical conditions, medication, tumour size, tumour grade, HR status and HER2 status and lymph node status are shown Supplementary Table 1a. The paired HV characteristics are shown in Supplementary Table 1b. The characteristics for patients with benign breast disease and paired HVs are in Supplementary Table 2. Patients with medical conditions or using medication which were known to influence neutrophil phenotype or function were excluded from the study (full list of inclusion and exclusion criteria can be found in Supplementary Table 3).

### Ethical considerations

Informed written consent was obtained from the patients and HVs on the day of blood taking itself, and samples were completely anonymised so participant identification could not be possible. The proposed human study was approved by Imperial Tissue bank committee and REC approval was obtained. It was also approved by the Crick ethical review committee for recruitment of HVs at the Crick. Material transfer agreements were signed by Imperial College and The Crick Institute to allow samples to be processed between the 2 sites. All work involving human participants was undertaken in accordance with Good Clinical Practice guidelines.

### Statistical considerations for human samples

Biological variability between individuals and size of effect calculations was estimated based on previous research data from investigating human neutrophil subpopulations^24^. Patients with EBC were selected based on HR status and HER2 receptor status. A control group of patients with benign breast disease and their paired HVs were also recruited. With the help of the Bioinformatics department at The Crick Institute, we estimated that ideally 21 patients were required for each subgroup of breast cancer patients and the group with benign breast disease and 11 paired healthy control volunteers (HVs) for each group were needed to obtain a statistical power at >80%, alpha = 0.05, to detect differences of 2.97 “units” in neutrophil subpopulations between groups (two-sided T-test). The method of Schouten was used to estimate the sample sizes under the assumption of unequal variances, sample sizes and independent samples, along with approximation to the t-test. Given the logistical challenges in recruiting patients during the COVID pandemic, we aimed to recruit 21 patients in each breast cancer subgroup but ended up recruiting fewer in each group.

Pamgene’s Bionavigator software (version 2.3, 2020) was used for image quantification and statistical analysis. This included a quality check to exclude any arrays that showed clear visual defects. Data was log-transformed and normalised to take into consideration any variation between patient and HV kinase samples related to neutrophil isolation on different experiment days. PCA was performed on the transformed data. A prediction of kinases responsible for the peptide phosphorylation changes were calculated using the Upstream Kinase Analysis tool in Bionavigator software. The Median Final score is calculated by combining a sensitivity score (difference between patients and HV groups) and specificity score (a set of peptides to kinase relationship which is predicted from 6 databases that Pamgene uses). The Median Kinase Statistic represents the differences between the groups, with effect size (values) and direction >0 is activation and <0 is inhibition. Results were also visualised using a kinase phylogenetic tree created via the online Coral tool (http://phanstiel-lab.med.unc.edu/).

#### Prediction Model

The raw data for the prediction model are provided in Supplementary file 1 and the R code used in Supplementary file 2.

#### Code Availability

All analysis was performed using R version 4.1.3 and the and the R code used is provided in Supplementary file 2.

### Blood collection, neutrophil isolation and analysis

Human blood samples (40ml) were collected tubes from HVs at the Crick Institute and Imperial College with informed consent. Blood was collected in heparinised tubes for neutrophil separation with Histopaque 1119 for the flow cytometry experiments for the first 8 patients and paired HVs (P01-P08). EDTA tubes were used for blood collection for neutrophil separation with the immunomagnetic neutrophil isolation kits (which was used for all Pamgene assays and flow cytometry experiments for patients P09-P44, benign patients and paired HVs). Blood was sent in an EDTA tube for Full Blood Count analysis at Charing Cross Hospital from patient P07 and paired HV onwards to calculate the NLR.

#### Neutrophil isolation

Density gradient centrifugation was used to isolate neutrophils from blood using Histopaque 1119 (pre-warmed to 37°C) followed by further centrifugation using Percoll Plus GE Healthcare. Neutrophils were resuspended in buffer (HBSS -Mg -Cl -Phenol red, 10Mm HEPES buffer, 0.1% human serum).

For the Kinome assay, neutrophil Isolation kits by Millentyi and Stem Cell technologies were used to isolate neutrophils from the blood which worked based on immunomagnetic negative selection. For patients P01-P14 and B01-B02 and the paired HVs, the Milltenyi neutrophil isolation kit was used (Catalogue number 130-104-434). However, the Stem Cell neutrophil isolation kit (Catalogue number 19666) was used for P15-P44 and B3-B9 and the respective paired HVs as this kit gave a much higher yield of neutrophils.

#### FACS analysis

Single cell suspensions from human blood (2.5 X10^6^ cells/tube) were incubated with 5% human FcR Blocking Reagent (Miltenyi) for 10 minutes on ice and then incubated for 30 minutes over ice with various specified fluorescent surface marker antibodies (including CD16, CD66b and CD62L (clones CLB-gran 11.5 (catalogue 746199), G10F5 (catalogue 564679) and DREG-56 (catalogue 740982) respectively, from BD Biosciences) using a 1:200 dilution). The LSRFortessa cell analyser running FACSDiva software (BD Biosciences) and FlowJo software was used.

#### Pamgene kinase assay

Neutrophils lysates were generated using the Pamgene standard manufacturer protocol and were stored at -80^0^C. Lysate samples were placed on STK Pamchips® and PTK Pamchips® and were run separately using the standard manufacturer protocol. Supplementary figures 4a and 4b represent an outline of the various steps involved for STK and PTK assays respectively. Images are taken every 5 minutes to generate kinetic data in real time which was analysed using Pamgene™ Bionagivator software.

### Plasma extraction

Blood was centrifuged at 300rcf for 10 minutes at 37°C. The yellow supernatant was collected and centrifuged at 2000rcf for 10 minutes. The supernatant was collected for the functional use immediately or snap frozen and stored at -80°C.

### Neutrophil half-life quantification using time-lapse imaging

Human neutrophils (5 x10^4^ cells/well) were seeded in a 96-well plate containing 10mM HEPES and 3% plasma (obtained from the cancer patient or paired HV). Then 0.2μM of Sytox Green (used to identify dead cells) and 4μg/ml of Hoechst (used to identify DNA and cells) were added to each well. Cells were taken to the inverted Nikon wide-field microscope system and measurements were recorded per well every 30 mins for 20-32 hours using a 40x objective. Quantification of Sytox positive cells compared with the total cells was performed using Fiji software. Half-life values represent the timepoint at which 50% of the cells within the field of view are Sytox positive.

## Data availability statement

Access to the minimum dataset that is needed to interpret, verify and extend research is provided in this paper and the supplementary material.

## Supporting information

Raw data used for the analysis

algorithm scripts

## Acknowledgements

We are grateful for the support from the Flow Cytometry Unit and the Biological Resources Unit at the Francis Crick Institute. We thank the Imperial Breast Cancer Research Team (especially Bana Ambasager, Christina Ma, Jessica Lin and Kelly Gleeson) for their dedication to recruiting patients and HVs from Charing Cross hospital. We thank the Imperial BRC, CRUK Centre and the Imperial ECMC for support. Thanks to Gita Mistry who helped recruit HVs from the Francis Crick institute. Many thanks to patients and HVs from Imperial College London NHS trust and the Francis Crick who donated blood samples for the study. We thank Professor Edwin Chilvers for his advice on the clinical study set up. Thanks to Savi Rangarajan from Pamgene company for guidance on using the Pamgene Bionavigator software to interpret the raw kinase data.

## Author contributions

AR helped with the study design, conducted most of the experiments, analysed data and helped interpret the results. FP performed animal experiments and analysed data. BA helped recruit patients and HVs and coordinated transferring blood samples and KG helped manage the team of Breast Cancer Research nurses. IVA conducted the ex-vitro culture experiments and analysed data. CL designed prediction model from neutrophil kinase activity. VP supervised the ex-vitro functional experiments. IM and RCC helped design and supervised the study, interpreted the data, and wrote the manuscript. AR, IVA and VP critically reviewed the manuscript.

---

**Supplementary Figure 1.**
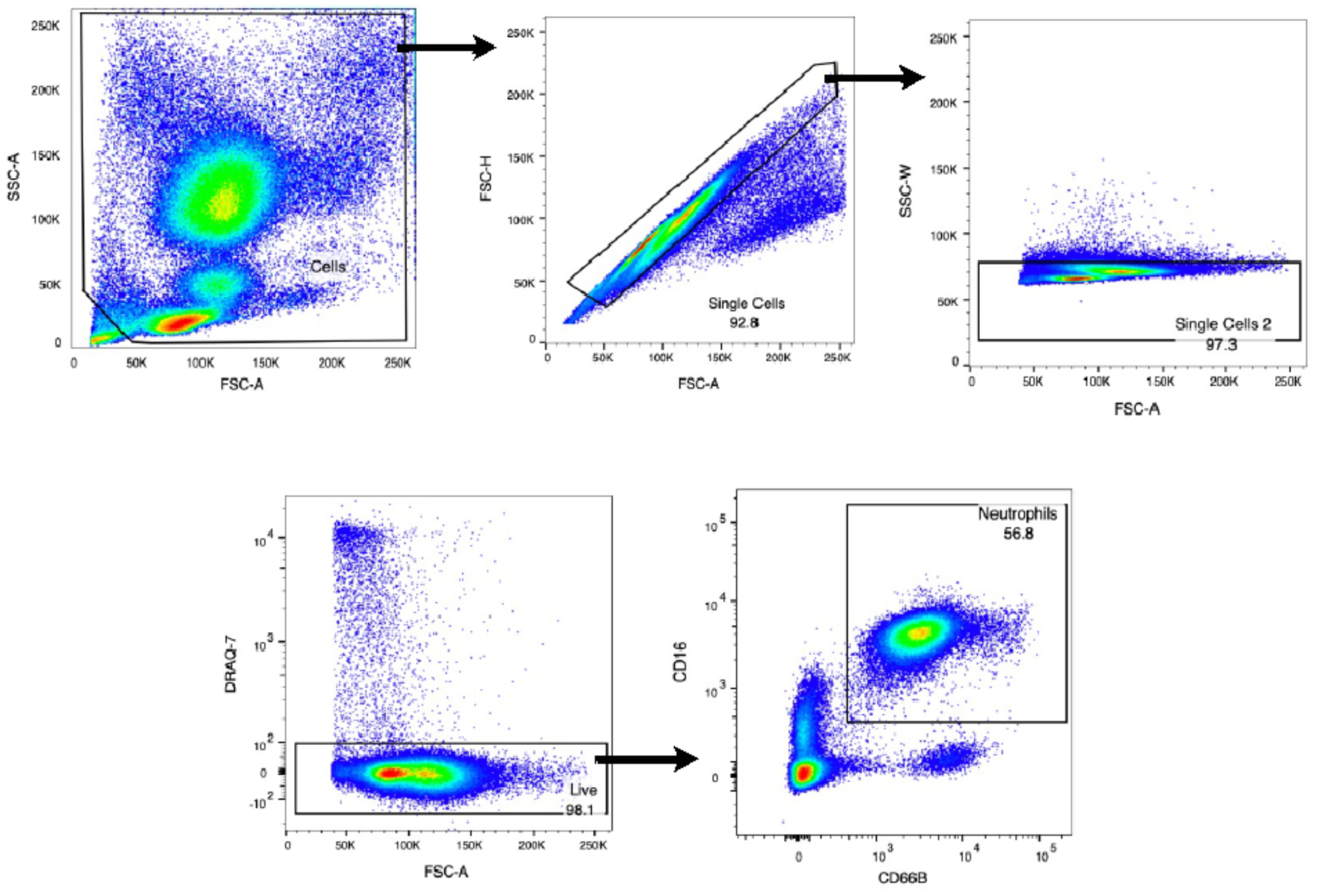
Flow cytometry strategy for human circulatory neutrophils (CD16+CD66B+).

**Supplementary Figure 2.**
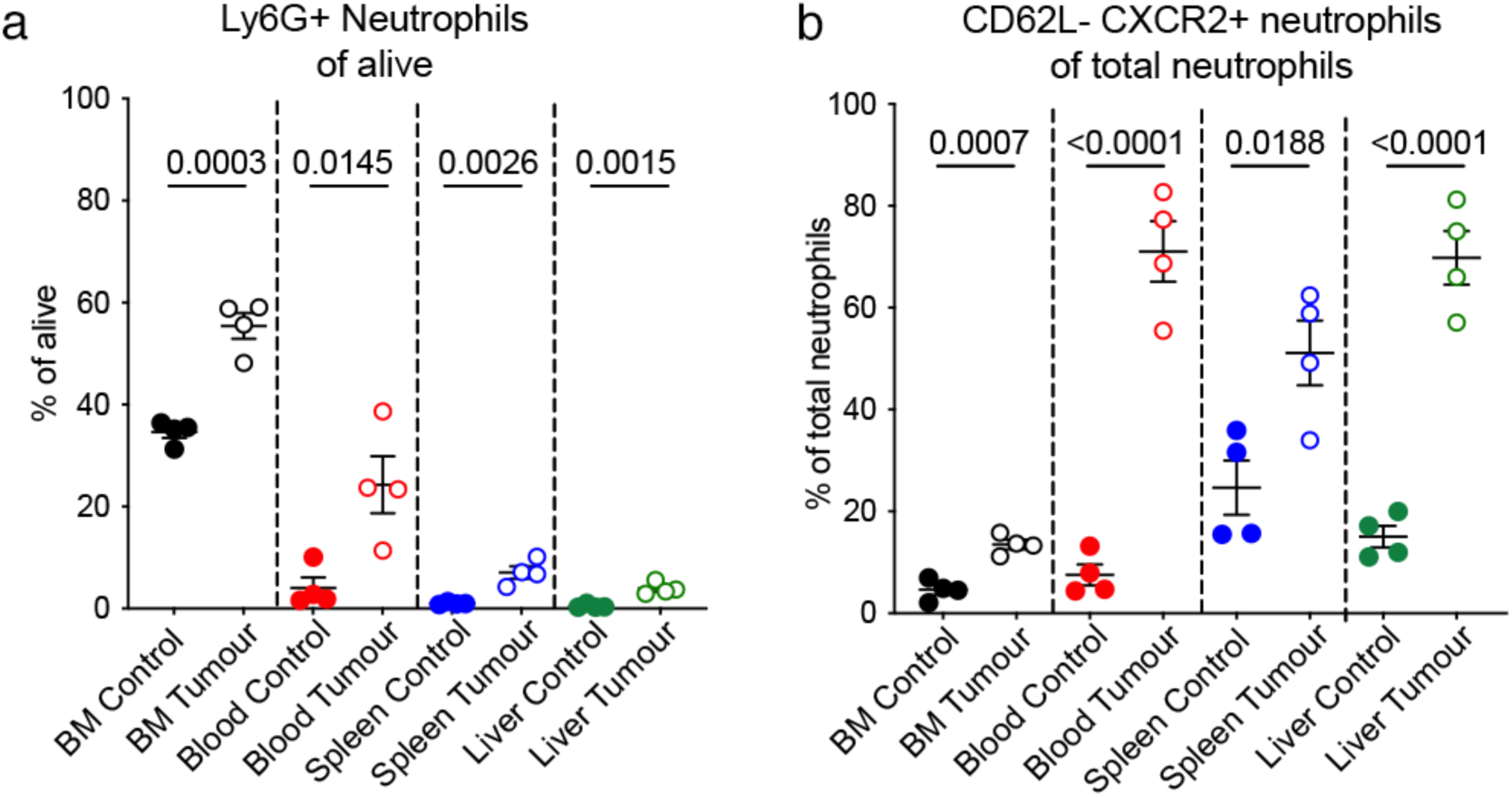
**a.** Flow cytometry showing neutrophils (LY6G^+^ CD11b^+^) in the bone marrow (BM), blood, spleen, liver and lungs of tumour-bearing mice and healthy controls. Each dot represents one mouse. Data are shown as mean ± standard deviation (SD). Statistical analysis by two-sided t-test. **b**. Flow cytometry showing proportion of CD62L-CXCR2+ of total neutrophils (LY6G^+^CD11b^+^) in the BM, blood, spleen and liver of tumour-bearing mice and healthy controls. Each dot represents one mouse. Data are shown as mean ± standard deviation (SD). Statistical analysis by two-sided t-test.

**Supplementary Figure 3.**
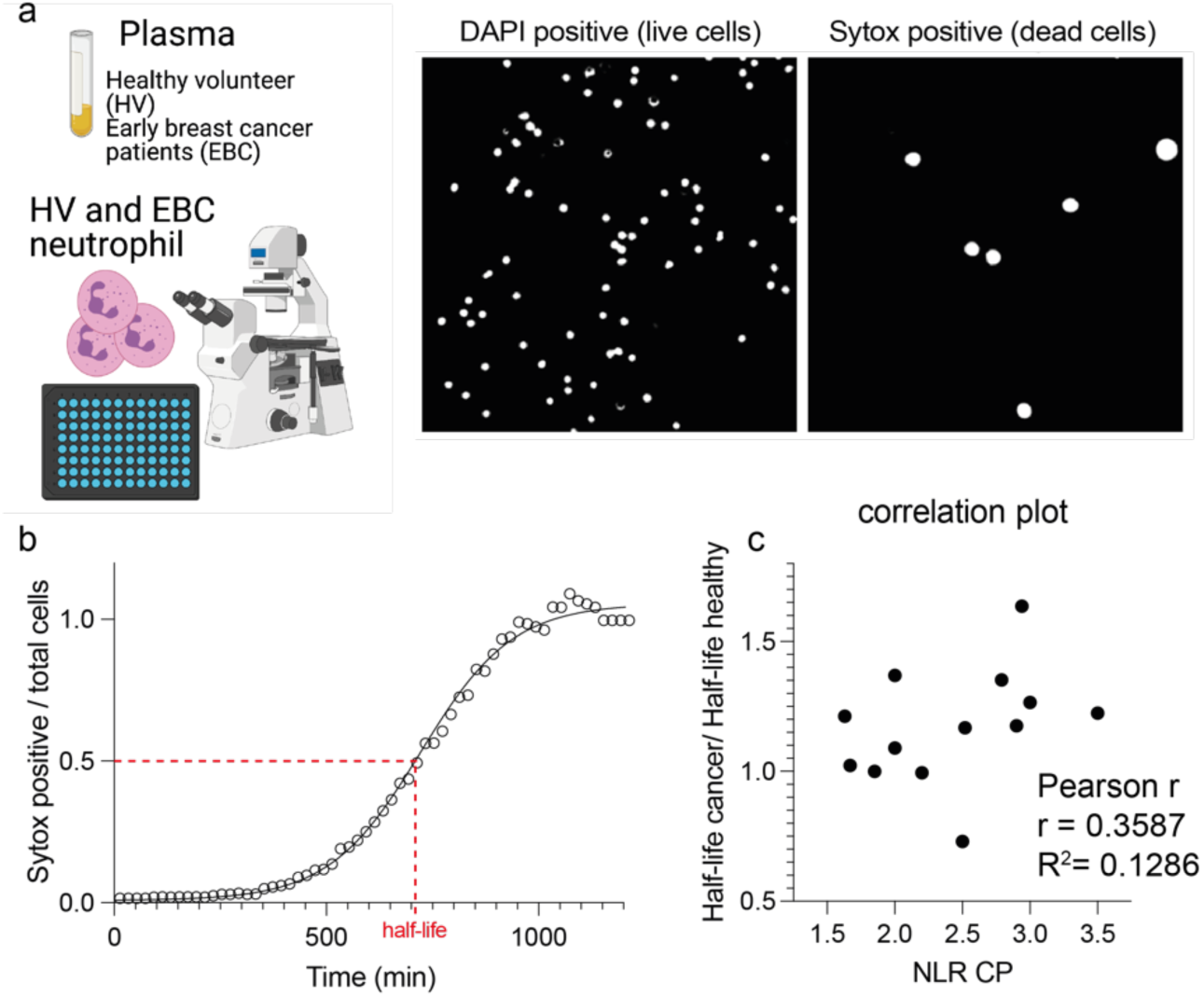
**a**. Scheme of the experimental set-up for the live cell imaging and half-life quantification. Example of live and dead cells identified by intensity-based thresholding at a specific timepoint. **b**. representative live curve of number of dead cells over total cells of a specific field of view over time and the half-live (dotted red line) representing the time at which 50% of the total cells are dead. **c**. Correlation analysis of the neutrophil to lymphocyte ratio measured in EBC patients and the fold change in the corresponding half-life value compared to the one obtained for the matching HV.

**Supplementary Figure 4.**
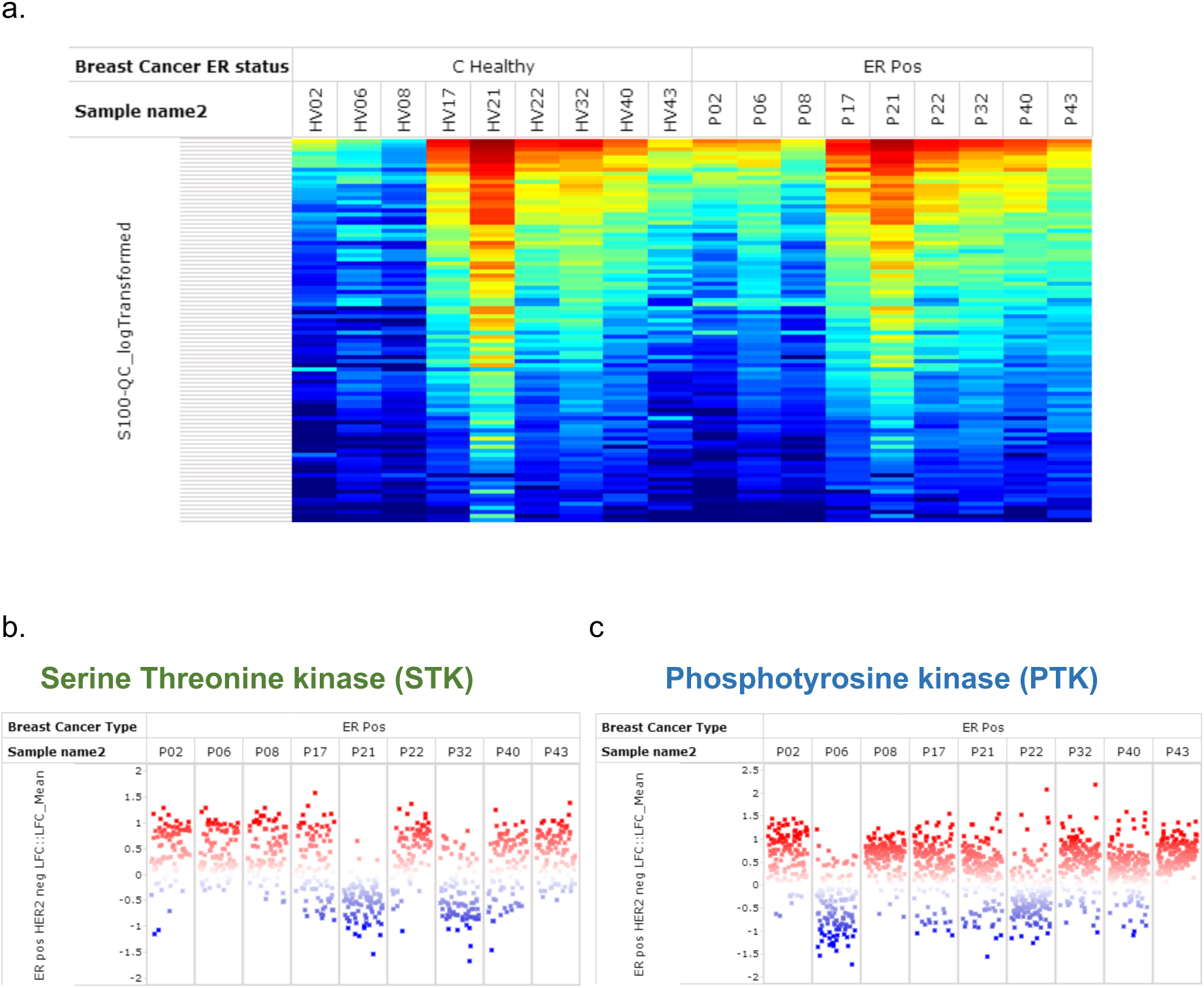
**a. Heatmap to show an overview of peptides which have been phosphorylated by kinases within the neutrophil lysates in HVs (represented by column C Healthy) and patients with HR-positive HER2 negative breast cancer (represented by ER pos column).** The sample name 2 row refers to the individual patients (letter P in front of numbers) and paired HV in front of numbers. Data represented prior to normalisation to accommodate for variability due to experiments done on different days **b-c Log-fold change (LFC) in phosphorylated peptides in patients with breast cancer compared to paired HVs (individual patients are represented by different columns) for STK (b) and PTK (c) family of kinases**

**Supplementary Figure 5.**
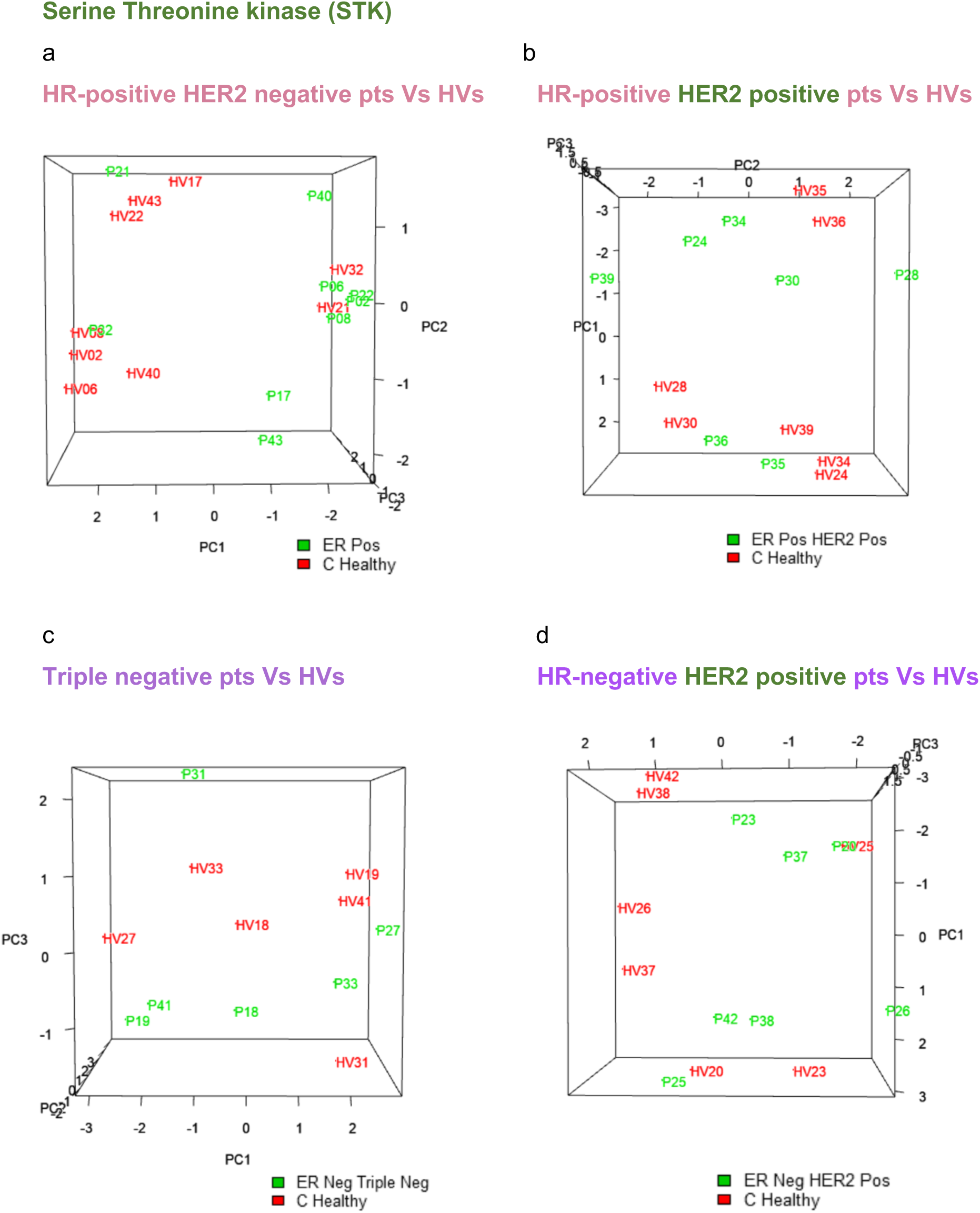
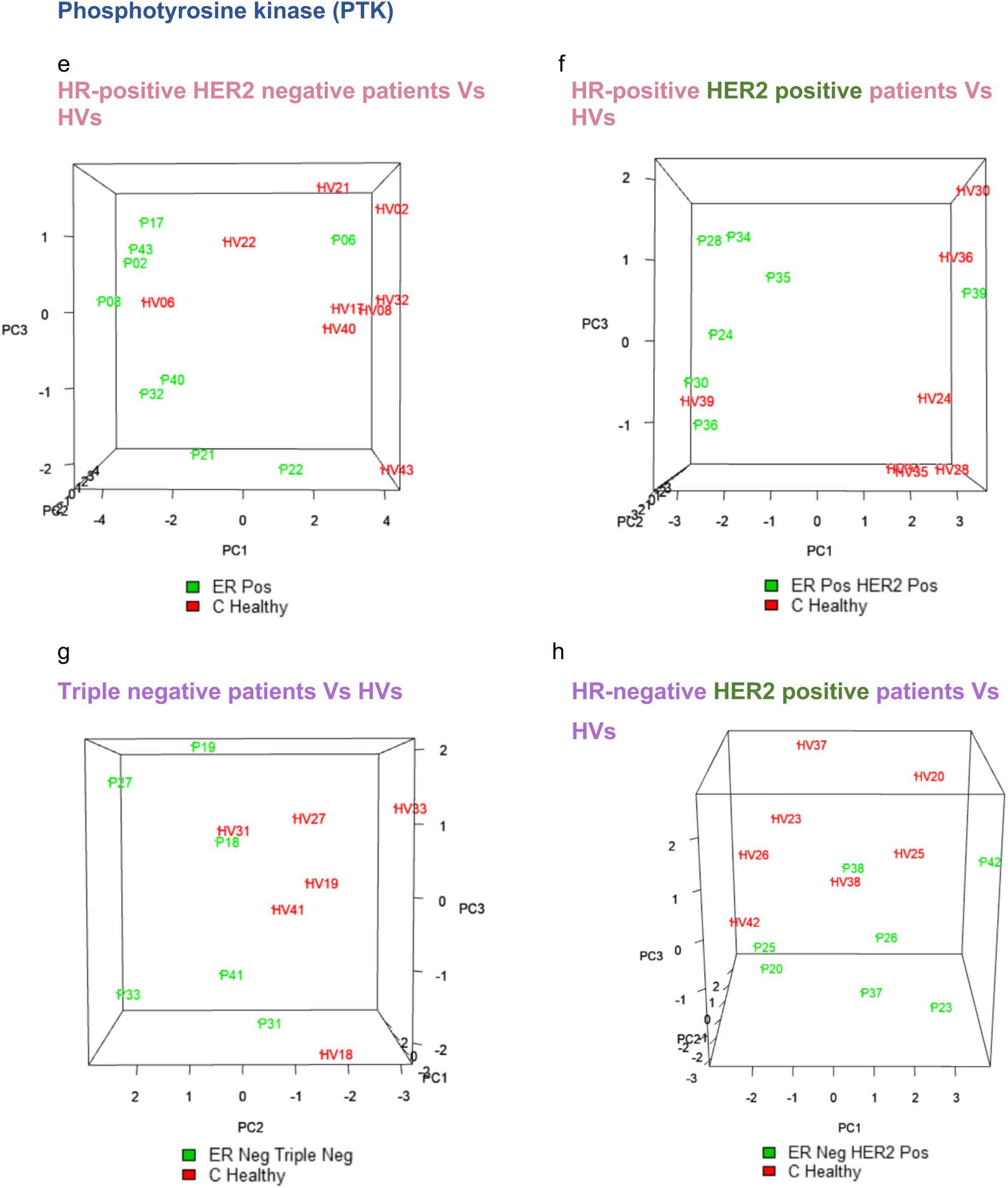
**a-d: Principal Component Analysis (PCA) of the different subtypes of breast cancer compared to HVs for the STK family of kinases.** a. HR-positive HER2 negative breast cancer (represented by ER pos) and HVs (represented by C Healthy); b: patients with HR-positive HER2 positive breast cancer (represented by ER pos HER2 pos) and HVs (represented by C Healthy); c: patients with HR-negative HER2 positive breast cancer (represented by ER neg HER2 pos) and HVs (represented by C Healthy); d: patients with Triple negative breast cancer (represented by Triple neg) and HVs (represented by C Healthy). **e-h Principal Component Analysis (PCA) of the different subtypes of breast cancer compared to HVs for the PTK family of kinases.** e: patients with HR-positive HER2 negative breast cancer (represented by ER pos) and HVs (represented by C Healthy); f: patients with HR-positive HER2 positive breast cancer (represented by ER pos HER2 pos) and HVs (represented by C Healthy); g: patients with HR-negative HER2 positive breast cancer (represented by ER neg HER2 pos) and HVs (represented by C Healthy); h: patients with Triple negative breast cancer (represented by Triple neg) and HVs (represented by C Healthy).

**Supplementary Figure 6.**
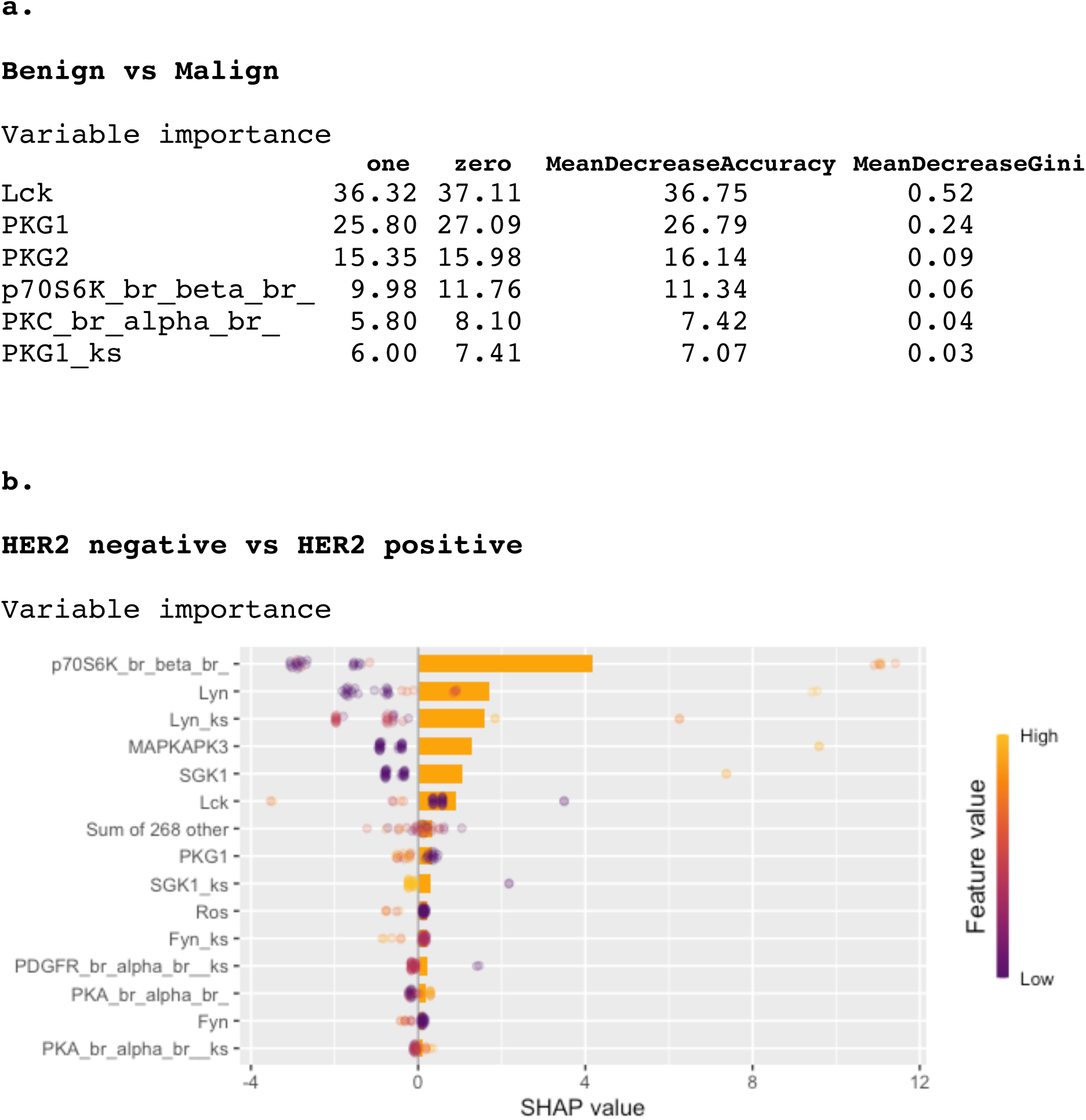

**Supplementary Figure 7.**
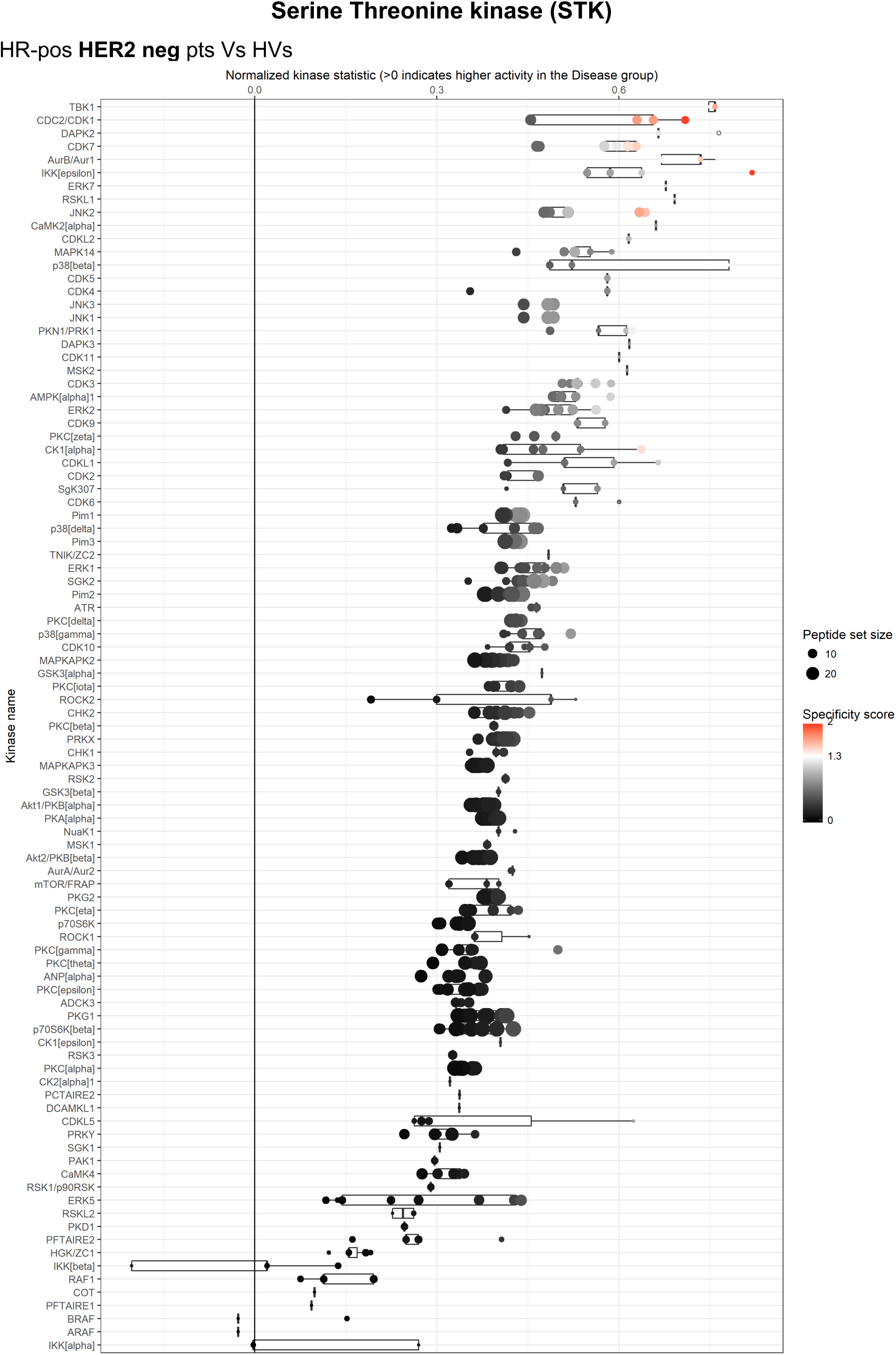
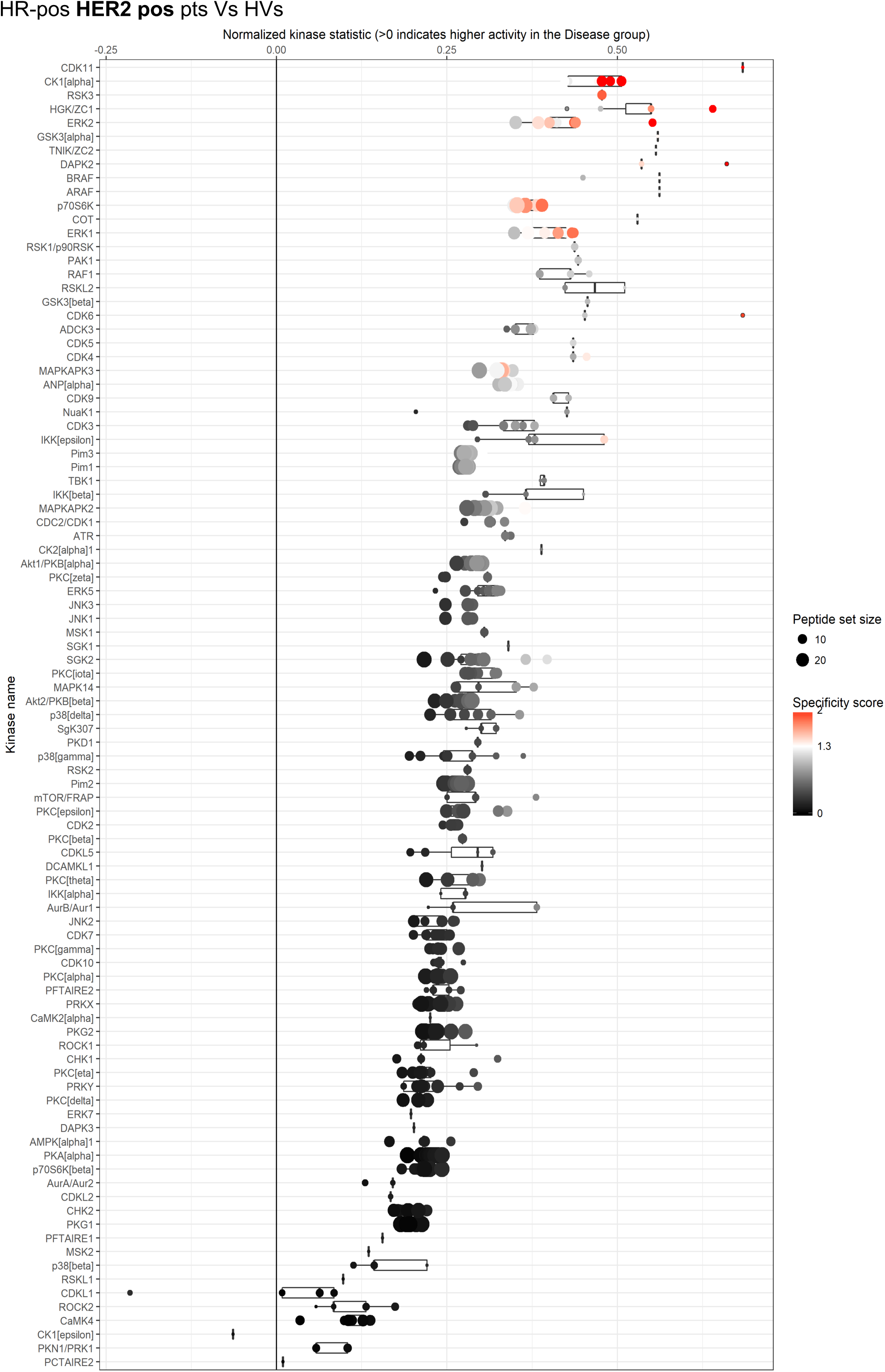
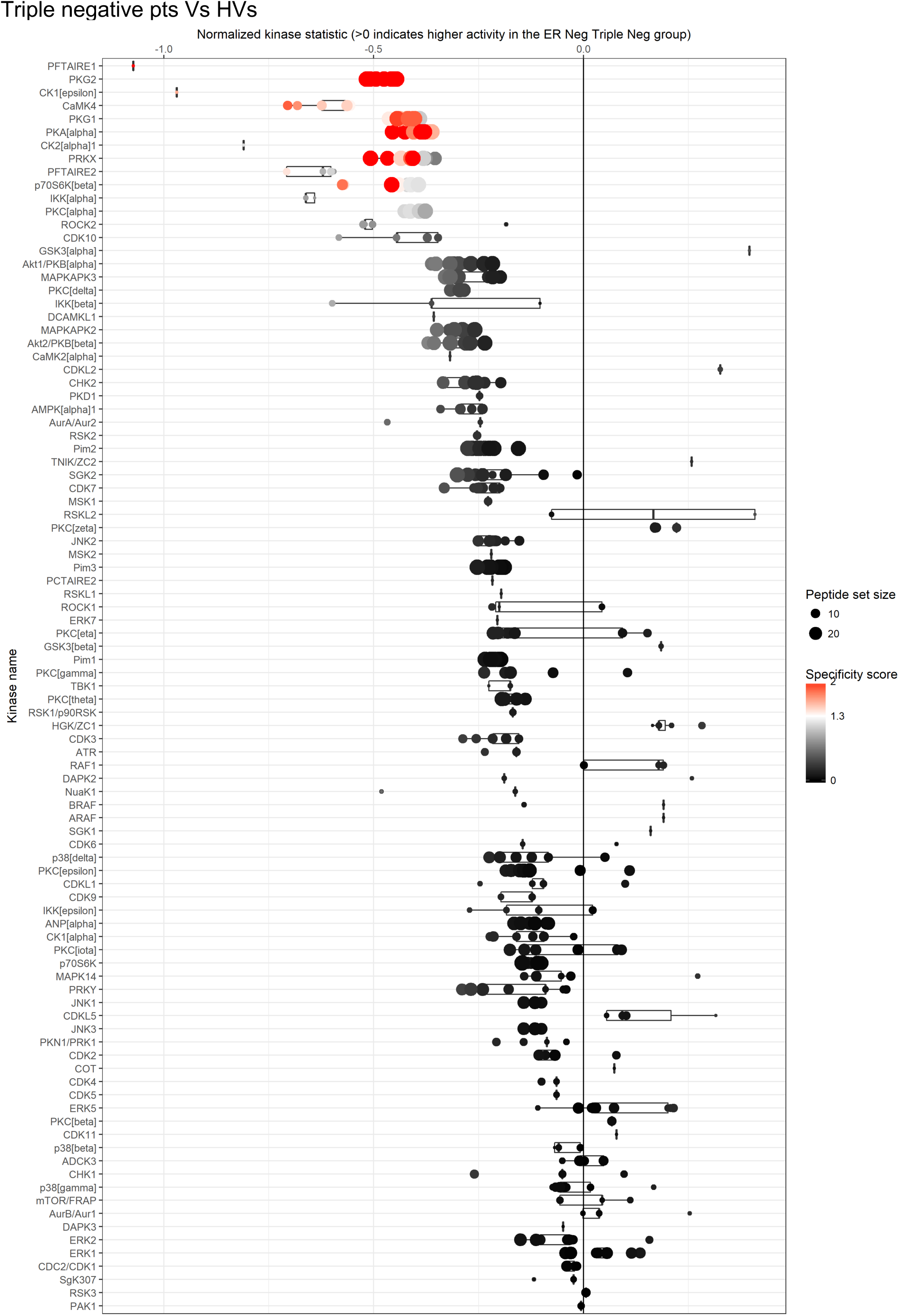
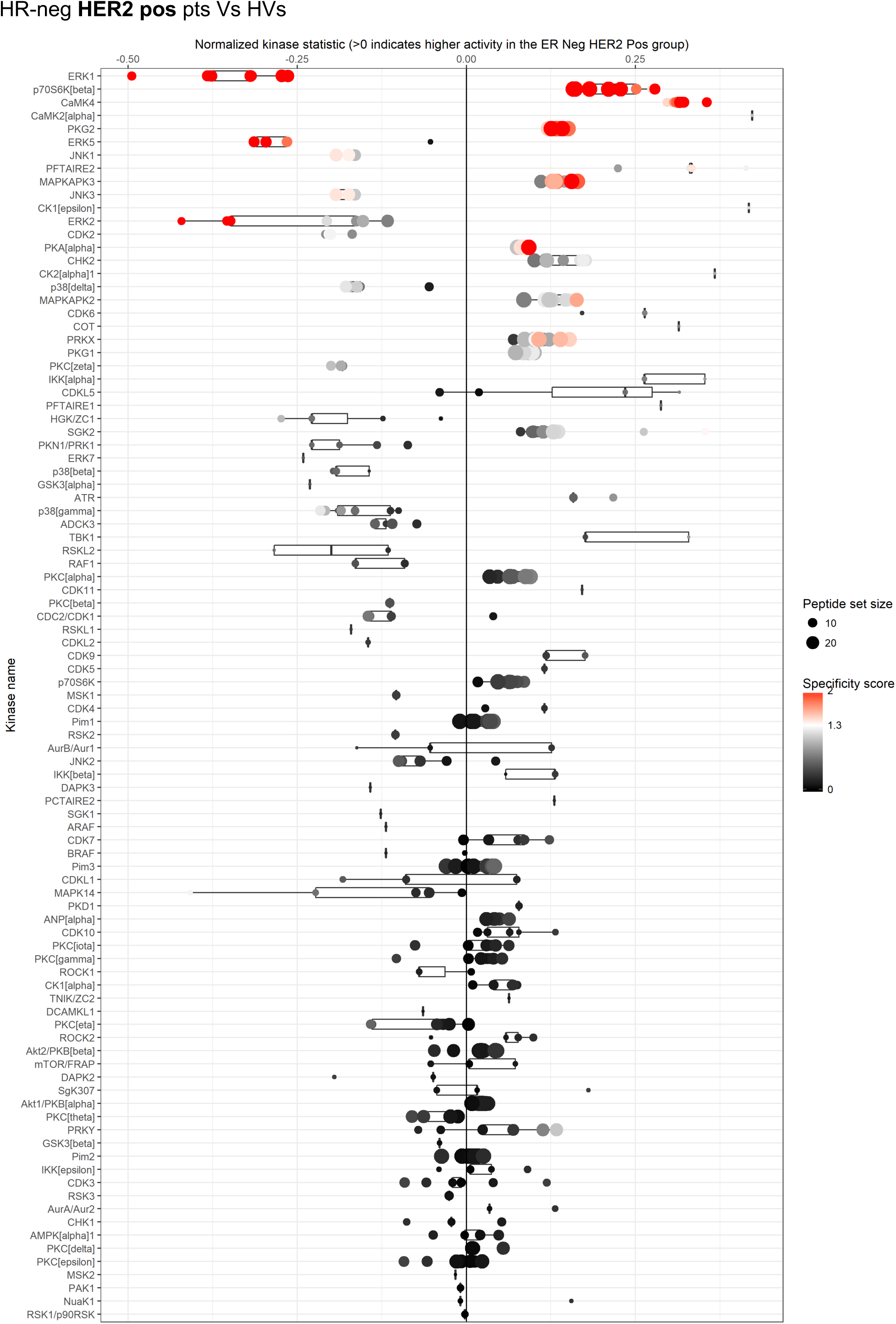
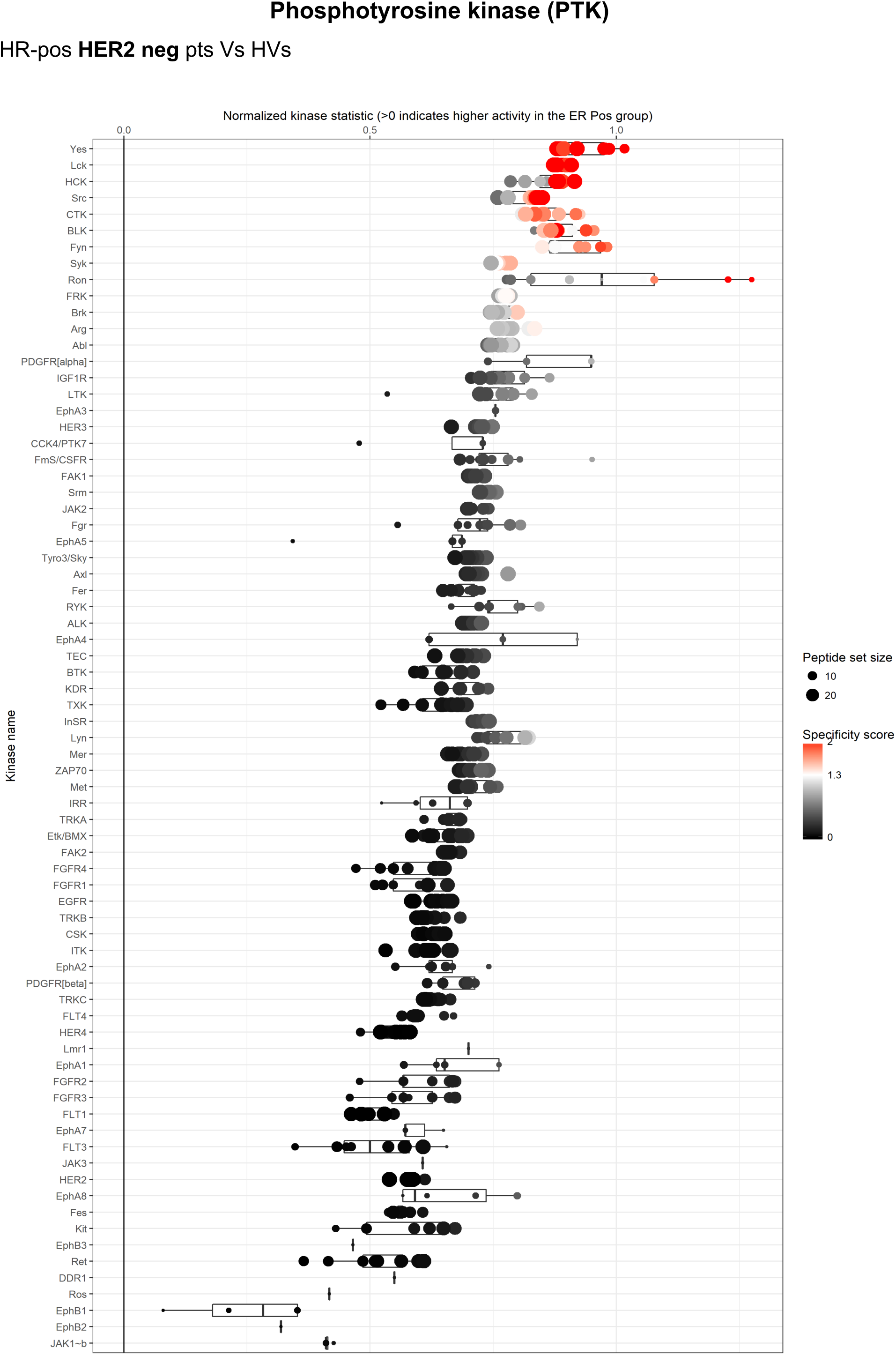
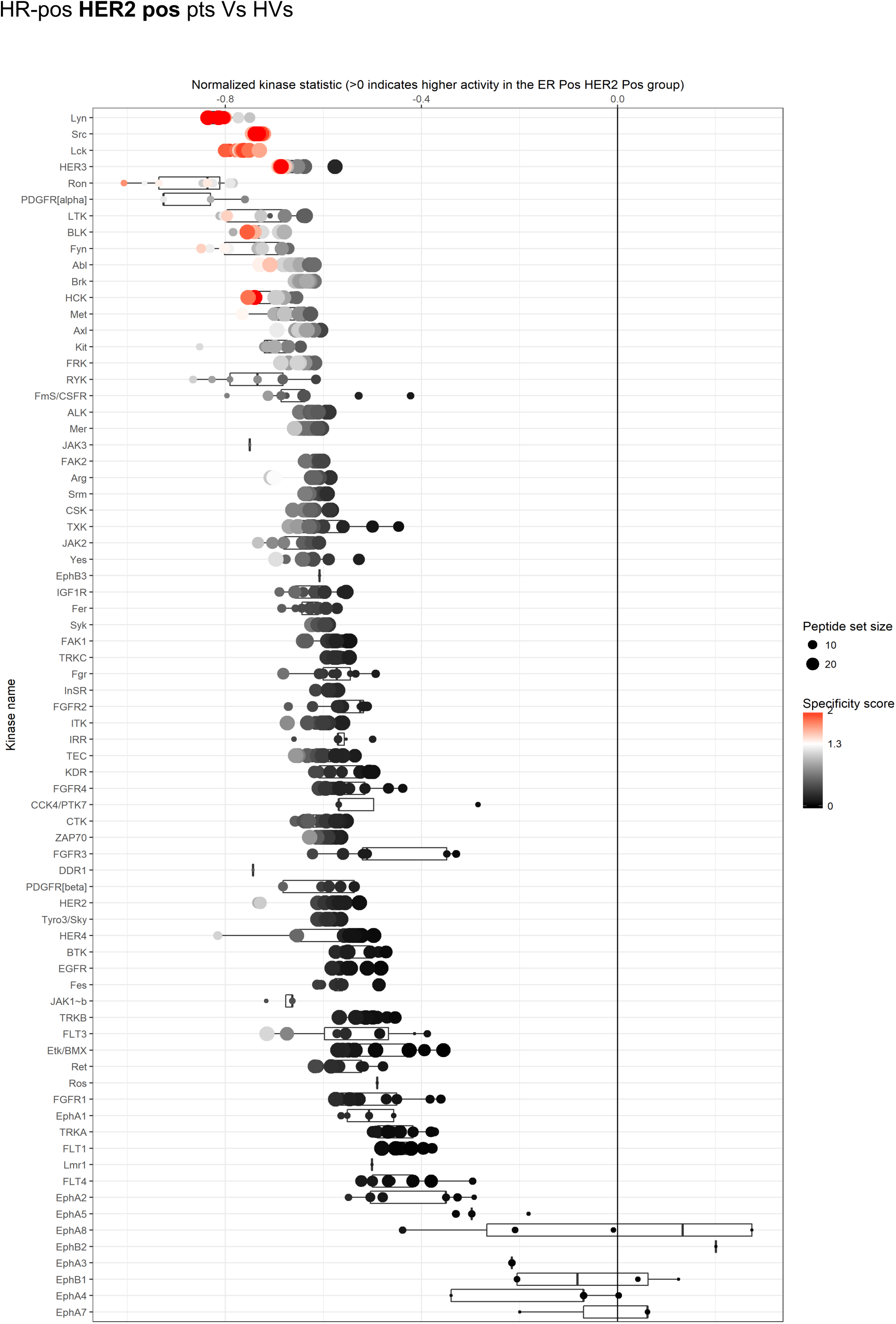
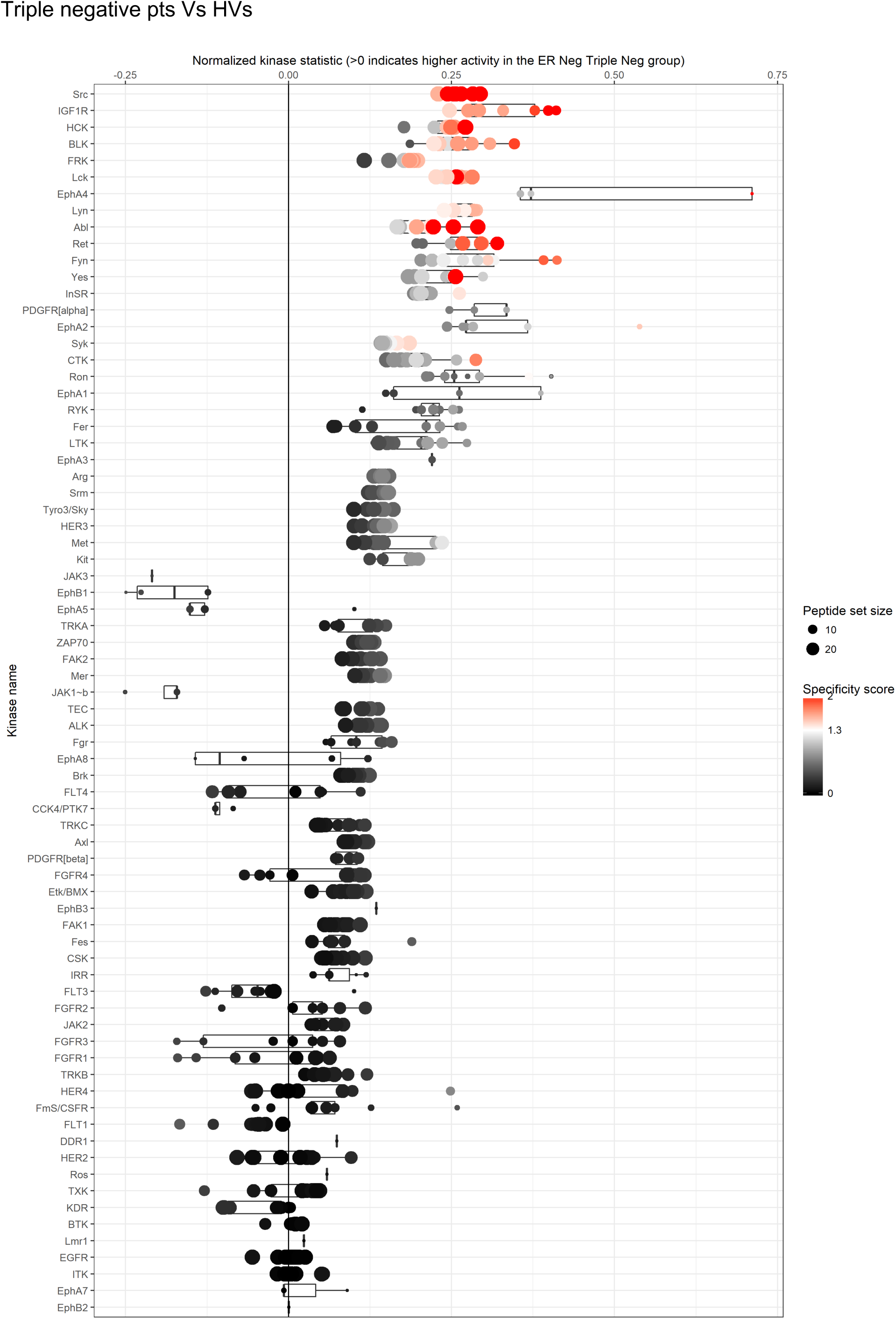
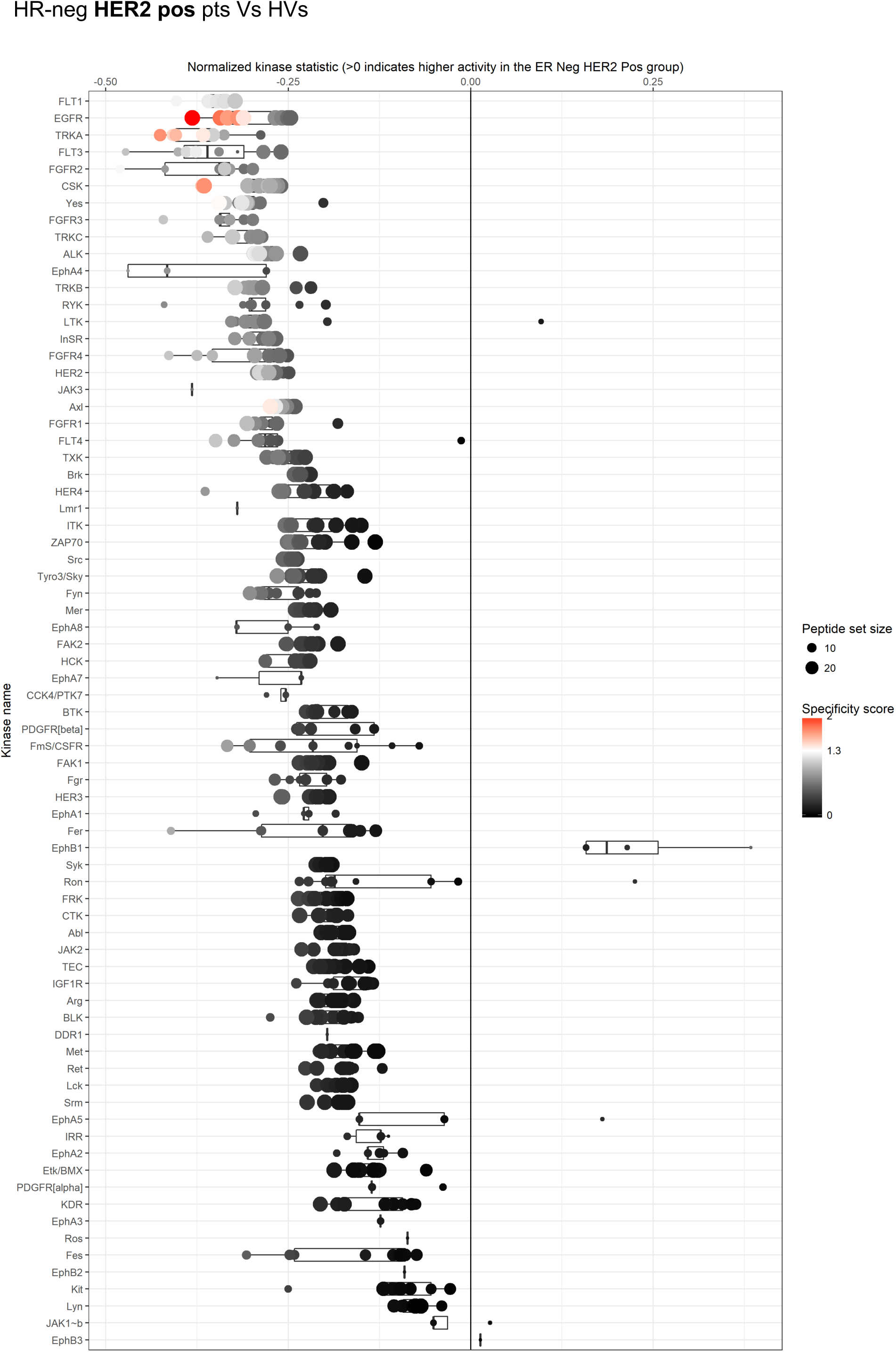
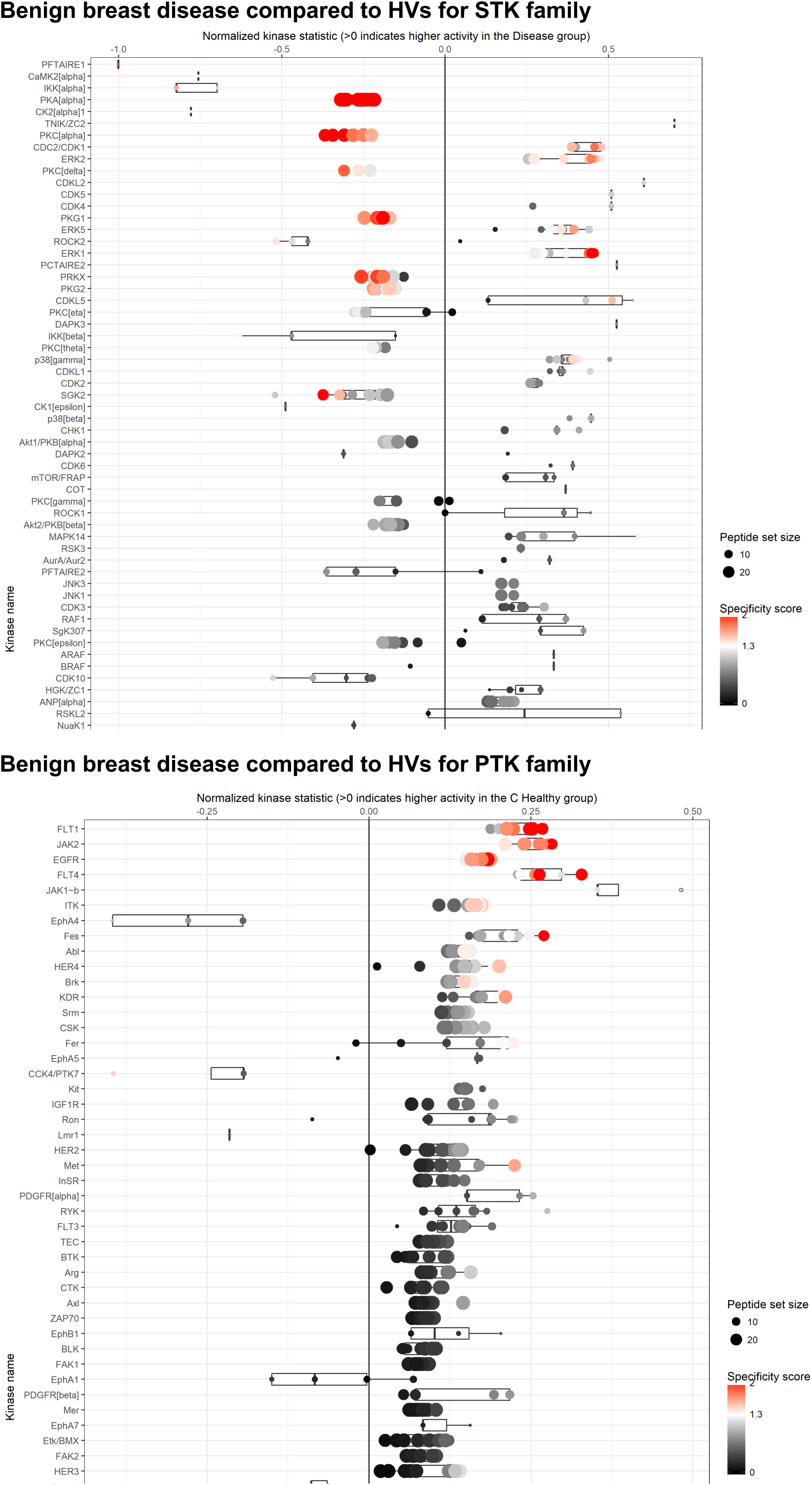
Upstream Kinase Analysis showing a list of putative kinases for either the STK or PTK families based on the peptides phosphorylated differently in patients compared to HVs. The analysis is done for each subtype of breast cancer or tpatients with benign disease separately as indicated. Size of circle represents peptide data set belonging to each kinase and colour scale represents the specificity of the kinase for that peptide set (red highly specific).

